# PanSpace: Fast and Scalable Indexing for Massive Bacterial Databases

**DOI:** 10.1101/2025.03.19.644115

**Authors:** Jorge Avila Cartes, Simone Ciccolella, Luca Denti, Raghuram Dandinasivara, Gianluca Della Vedova, Paola Bonizzoni, Alexander Schönhuth

## Abstract

**Motivation:** Species identification is a critical task in agriculture, food processing, and health-care. The rapid growth of genomic databases — driven in part by the increasing investigation of bacterial genomes in clinical microbiology — has outpaced the capabilities of conventional tools such as BLAST for basic search and query tasks. A key bottleneck in microbiome studies lies in building indexes that allow rapid species identification and classification from assemblies while scaling efficiently to massive resources such as the AllTheBacteria database, thus enabling large-scale analyses to be performed even on a common laptop.

**Results:** We introduce `PanSpace`, the first convolutional neural network–based approach that leverages dense vector (embedding) indexing —– scalable to billions of embeddings —– for indexing and querying massive bacterial genome databases. `PanSpace` is specifically designed to classify bacterial draft assemblies. Compared to the most recent and competitive tool for this task, `PanSpace` requires only ~2 GB of disk space to index the AllTheBacteria database, an 8*×* reduction relative to existing methods. Moreover, it delivers ultra-fast query performance, processing more than 1,000 assemblies in less than two and a half minutes, while preserving the utmost accuracy of state-of-the-art approaches.

**Availability:** `PanSpace` is available at https://github.com/pg-space/panspace.

## 1 Introduction

Determining the species a DNA fragment stems from is one of the most fundamental tasks in biology. BLAST [1], one of the most popular, if not the best known tool in bioinformatics, addresses this problem. BLAST queries a database storing large collections of annotated genomes by aligning sequenced DNA against the genomes of the collection. Various fine-tuned heuristics are necessary to guarantee that those queries are answered in reasonable amounts of time. Upon having queried the database, one can analyze the resulting highest-scoring alignments to determine the evolutionary origin of the query fragment.

In bacteria, the first definition of *species* was based on DNA-DNA hybridization (DDH). It was found reasonable to classify two strains to belong to the same species if their DNA hybridized at at least 70% [47]. In the following, the 70% threshold was routinely used as an operational demarcation in DDH based species classification. However, DDH is time-consuming and labor-intensive. As a consequence, alternative, less expensive experimental protocols were in great demand. The steadily increasing number of sequenced genomes further aggravated the impracticability of DDH. The finding that Average Nucleotide Identity (ANI) matched DDH as a similarly reliable basis to distinguish between the evolutionary origin of DNA fragments paved the way for algorithmic approaches. In the meantime, algorithms that address the computation of the ANI have become widely popular for delineating between species. These algorithms both evaluate the output of whole genome and gene based protocols [12, 16, 26, 35]. In particular, these studies indicate that a DDH threshold of 70% corresponds to an ANI threshold of approximately 95-96% [12].

Genomic databases are becoming increasingly larger, now storing millions of genomes — NCBI RefSeq counts 2.3 million genomes [14, 37], GISAID 20 millions^1^, and ENA 1.9 millions [15]. However, querying DNA sequence databases with alignment-based search tools requires substantial computational resources due to the algorithmic complexity of the corresponding alignment problems. In fact, running BLAST on any of those databases proves impossible if not done on gigantic compute clusters.

In summary, when ordinary research laboratories look to classify DNA sequences in terms of their evolutionary origin, they have to resort to alignment-free approaches. In this paper, we will present such an alignment-free approach, which to the best of our knowledge for the first time is genuinely artificial intelligence (AI) based. While highly accurate alignment-free approaches have already been existing in the meantime, the decisive achievement of our approach is its resource friendliness: its RAM and runtime requirements are reduced by up to orders of magnitude in comparison with the state of the art. Importantly, our approach preserves the great classification accuracy of the prior approaches.

In somewhat more detail, we focus on the classification of prokaryotes at the taxonomic level of species. Seminal work [49, 50] significantly advanced the in the meantime large field of bacterial taxonomies. In that initial work, phylogenetic trees referring to only the 16S rRNA gene were constructed. The choice of the 16S rRNA gene, known as the “subunit rRNA gene”, is motivated by its high level of conservation and its omnipresence in prokaryotes [43]. Another reason for its choice when identifying bacterial species [33, `TYGS`] is its short length (ca. 1485 bp) which supports rapid pairwise algorithmic comparisons.

The non-negligible drawback of 16S rRNA gene based approaches is the potential absence of this gene when working with draft shotgun assemblies. Because such assemblies have become standard in the analysis of prokaryotes, the 16S rRNA gene is no longer a reliable basis for exploring the evolutionary spaces of prokaryotes. Moreover, the short length, labeled above as a decisive advantage, can turn into a decisive disadvantage when working with databases that store not just thousands, but millions of prokaryotic genomes, because the possible combinations of species-specific mutations harbored by the 16S rRNA gene are limited. Resorting to highly conserved genes of greater length, on the other hand, prevents the computation of sufficiently accurate alignments because of the computational complexity inherent to the alignment problem.

This analysis of current experimental and algorithmic opportunities explains why alignment-free approaches are currently obviously the most advantageous and promising ones. In the meantime, various such approaches have been presented; see [24, `fIDBAC`] and [29, `GAMBIT`], for example. In particular, all of these approaches are rooted in the evaluation of the composition of *k*-mers of the genomes one compares, where *k*-mers are the subsequences of length *k* of the genomes, with *k* being relatively short (e.g. *k* = 4, 6, 8).

Importantly, any such approach requires a curated database that hosts thousands of genomes as a reference. In this, the rapid growth of such databases opens up opportunities to analyze larger sets of sequences from the point of view of pangenomes, that is, by making use of advanced data structures that support the referencing of entire populations of sequences. In the following, we list some of the most relevant databases that serve these purposes. The Type Genome Server database (`TYGS`)^2^ [33] contains more than 20 000 microbial-type strain genomes, which are updated weekly [2, 7]. `TYGS` incorporates quality checks to reject poor quality assemblies, and incorrectly identified species. `fIDBAC` [24] stores assemblies of bacterial genomes, together with metadata that specifies the corresponding strains. `fIDBAC` uses CheckM [34] to evaluate the completeness and contamination of each genome. Some additional curation steps that involve the information provided by the LTP [53] database and information on strains from LPSN [53], allow to discard mistakenly labeled genomes. In an overall account, the `fIDBAC` database contains more than 12 000 bacterial genome assemblies, covering 9 827 species belonging to 2 448 genera. When referencing `fIDBAC`, 16S sequences are aligned against the sequences stored in the LTP database, while *k*-mers are matched with the *k*-mers in `fIDBAC` using KmerFinder[6]. `GAMBIT` [29] is a curated database that lists 48 000 bacterial genomes from the NCBI RefSeq database that span 1 414 species and 454 unique genera. To perform species identification, an approximate Jaccard similarity between the query and the *k*-mers of the genomes in the database is computed.

The current state-of-the-art for bacterial genome classification is `GSearch` [56]. `GSearch` is an alignment-free genome search tool that can work at pangenome scale, combining memory-efficient sketching algorithms such as MinHash [5] or HyperLogLog [11] for genomic distance estimation and Hierarchical Navigable Small World graphs (HNSW) [30] for finding nearest neighbors and identifying the originating species. A subset of the *k*-mers of the query genome and its occurrences is selected as a feature representation of each assembly and an index is created based on HNSW. The index is then queried using a Jaccard-like similarity of their feature representation.

Most recently, the largest available bacterial dataset, AllTheBacteria [15], was released. AllTheBacteria comprises all known bacterial genomes (more than 1.9 million draft assemblies), available from the the European Nucleotide Archive (ENA). Two alignment tools, `Phylign` [4] and `LexicMap` [40], make use of AllTheBacteria by first querying the database with a set of reads or contigs, and subsequently computing posterior alignments of the queried assemblies with the assemblies in the database suggested by the query as matches.

For querying, both `Phylign` and `LexicMap` construct k-mer-based indexes, where `Phylign` leverages a COBS index [3] of 248GB, while `LexicMap` extracts some *k*-mers for each genome that guarantee matches in short windows, which results in an index of 2.46TB. Subsequent alignments are determined using minimap2 [23] in the case of `Phylign`, and with the wavefront alignment algorithm [31] in the case of `LexicMap`. Even though classification is not the explicit purpose of those tools, one can naturally employ them for that purpose by aggregating the results of the query for each contig of a draft assembly.

Further but more distantly related approaches are tools that classify reads resulting from metagenome sequencing. Unlike in our case, such approaches operate on (raw) sequencing reads instead of draft assemblies. The goal of such approaches is to label each read in a metagenome sample, usually at the level of species. Prominent examples of such tools are `Kraken` [52], `Kraken2` [51], `Centrifuge`[20], and `Centrifuger`[42]. All of them focus on short reads, where `Centrifuger`, as the only exception, can also process long reads. Another tool that can handle long reads is `Taxor` [44].

To avoid dependency on databases, thereby decreasing the amount of computational resources required for classification, various machine learning tools have recently been suggested. Of course, because such tools circumvent classification of assemblies relative to the ultra-large databases available, they suffer from the corresponding drawbacks, which leads to reduced accuracy in terms of the predicted taxonomic information. Notable examples are `HiTaxon` [45] and `Taxometer` [21], both designed to work on bacterial genomes, while `DeepMicrobes` [25] specializes in the evaluation of gut microbiota. Tools of more limited scope are `EukRep` [48] and `Tiara` [19], both of which classify at the domain level, and `MycoAI` [36] which targets fungal sequences. While the major purpose of these tools is the classification of eukaryotic sequences, they have also been shown to work on bacterial data. An additional advantage of tools that work in immediate connection to databases is the fact that it is desirable in a wide range of applications not only to identify the taxon, but also reveal the sequences that provide evidence of the origin of the queries and that provably stem from the taxon in question. The reason is that these sequences support the performance of downstream analyses–importantly, index-based methods like ours preserve such qualities.

We propose `PanSpace`, a tool designed to accurately classify bacterial genomic sequences at the level of species. To this end, `PanSpace` integrates novel efficient approaches for both indexing and querying bacterial pangenomes. The major advantage and achievement of `PanSpace` is its economic usage of space and time. `PanSpace` indexes the AllTheBacteria dataset using less than 2GB of disk space and can process a thousand queries in less than two and a half minutes. `PanSpace` queries require less than 7GB of RAM for indexing and querying, making it possible to run `PanSpace` on a personal computer, without, for example, having to upload data to cloud servers.

We compare the results of `PanSpace` with `GSearch`, which is the current, clearly leading state-of-the-art approach. When comparing `PanSpace` with the most space-efficient version of `GSearch`, the `PanSpace` index is 8× smaller and requires 5.7× less RAM for its construction. When comparing `PanSpace` with the most accurate version of `GSearch`, the index of `PanSpace` is 47× smaller, and requires 21× less RAM for its construction, while preserving the high level of accuracy of `GSearch`.

### 1.1 Main Contributions

- We index the largest publicly available database of bacterial draft assemblies (the AllTheBacteria data set [15]), requiring approximately 2 GB of disk space, an 8-fold improvement over `GSearch`, which is the state-of-the-art.
- We experimentally show that the ANI between a query assembly and the assemblies resulting from querying our index exceed 95, which is the ANI that has been shown to reliably serve as a threshold when ascertaining that two assemblies belong to the same species.
- We propose the first embedding-based index for draft assemblies using the FAISS [18] index, with a Metric learning-based method that exploits FCGR representation of assemblies for the construction of these embeddings.
- We introduce the first machine-learning framework for prokaryotic genome classification that integrates index-based retrieval, providing both accurate taxonomic identification and direct access to the genomic evidence underlying each prediction.

## 2 Material and Methods

### 2.1 Overview of our approach

We propose `PanSpace`, a framework to index genomic sequences in a continuous *n*-dimensional space, referred to as *Embedding space*. Genomic sequences are represented by the Frequency matrix of the Chaos Game Representation of DNA (FCGR) [9], where the frequency of *k*-mers (4^*k*^ in total) are placed in a 2^*k*^ × 2^*k*^ matrix, where each cell correspond to a *k*-mer scaled frequency. In this representation, *k*-mers sharing an *l*-long suffix (with *l < k*) are placed contiguously in a 2^*k*−*l*^ × 2^*k*−*l*^ square matrix (see Figure 2). The FCGRs are used as feature representations of the sequences and then mapped to the Embedding space (*n* ≪ 4^*k*^) by leveraging a Convolutional Neural Network designed to exploit the *k*-mer ordering in the FCGR with Metric Learning for clustering based on species labels. The embedding representations are indexed with FAISS [18], an index for dense vectors, proven to scale up to 1 billion embeddings. The species identification step is done by querying the index and assigning the most common species among its closest neighbors w.r.t. the embedding representation of the assemblies.

Let us define by Σ = {*A, C, G, T* } the DNA alphabet, and the extended DNA alphabet by Σ_*N*_ = {*A, C, G, T, N*}, where *N* indicates a null call. Consider a collection 𝒮 of (possibly draft) assemblies, where each assembly *s* ∈ 𝒮 could be a set of more than one (contig) sequence over Σ_*N*_, and a set ℒ of species (labels). We assume there exists a labeling function *λ* : 𝒮 1→ ℒ that assigns a species in ℒ to each assembly *s* ∈ 𝒮 — learning such a function is the main goal of our method. The mapping 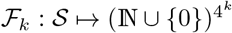 computes the ocurrences of all *k*-mers for an assembly.

### 2.2 Chaos Game Representation of DNA

Let us now introduce the key ingredients related to the FCGR representation of an assembly, which is the matrix representation of its *k*-mer occurrences mentioned in the previous section. We first define the Chaos Game Representation (CGR) of a DNA sequence *s* over the alphabet Σ. Observe that, as will be explained later, the FCGR k-mer representation is essentially a 2D-image (see Figure 1).

**Figure 1.**
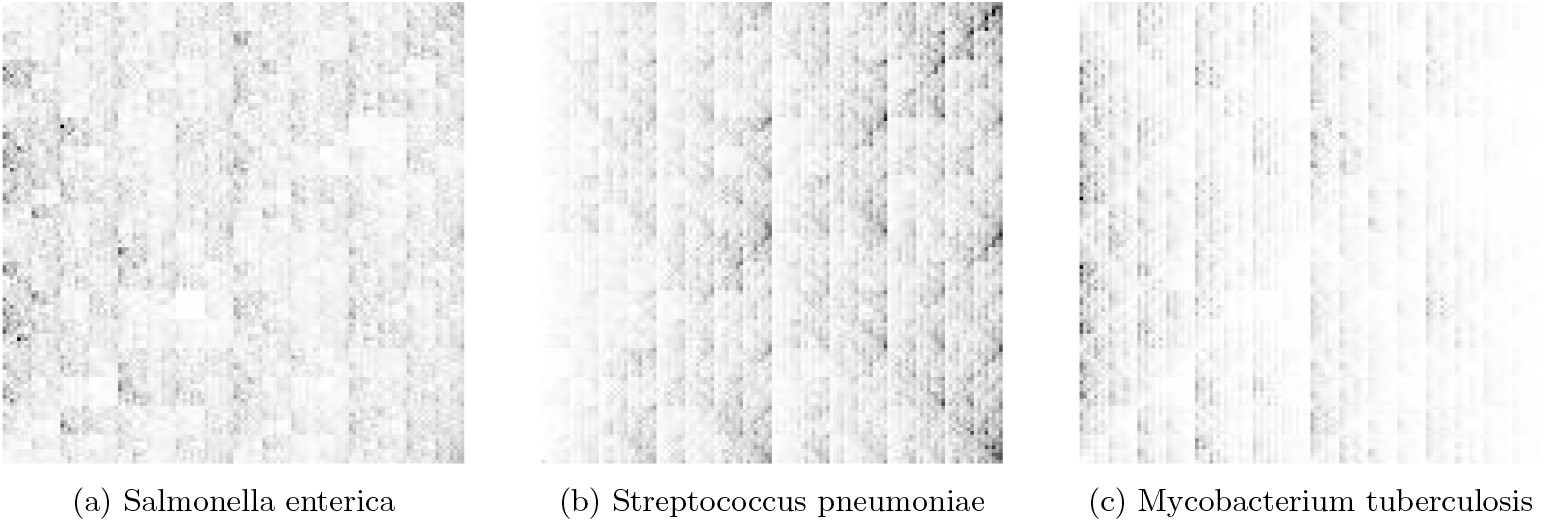
**Examples of FCGR** of assemblies from 3 species from the AllTheBacteria dataset. From left to right, Salmonella enterica (SAMD00003784), Streptococcus pneumoniae (SAMD00032020), and Mycobacterium tuberculosis (SAMD00111306). Each image corresponds to the visualization of the FCGR in 8-bits (*k*-mer frequencies rescaled to the interval [0, 255]) for *k* = 7. White color in a pixel means that the *k*-mer encoded has the minimum value of frequency found in the assembly. Black color represents *k*-mers with the maximum frequency in the assembly. Gray colors correspond to *k*-mer frequencies lying between the minimum and maximum values.

The CGR encoding of a DNA (and RNA) sequence is a coordinate in ℝ^2^, and is formally defined [17] as follows.

**Definition 1** (Chaos Game Representation of DNA (CGR)). *Given a sequence s* = *s*_1_ · · · *s*_*n*_ ∈ Σ^+^, *the CGR encoding of s is the ordered pair* (*x*_*n*_, *y*_*n*_) *which is defined iteratively as*

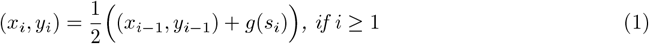

*where* (*x*_0_, *y*_0_) = (0, 0) *and*,

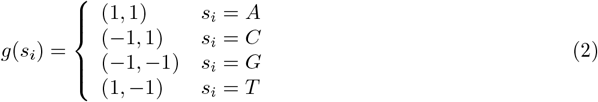

When plotting all the points resulting from iteratively encoding all characters in a DNA sequence, we obtain images like the ones shown in Figure 1. This requires an entire sequence over the alphabet Σ, and poses two challenges in real settings: sequences usually have missing data, *i*.*e*. they are defined over the alphabet Σ_*N*_, and they are often present as draft assemblies; as collections of contigs. The CGR encoding of a sequence is reversible, which means that each sequence has a unique 2-dimensional position in the square [1, −1], in particular, we can use it to find the positions of *k*-mers in the square (see Figure 2). As shown in [9, 46] the CGR visual representation is well approximated by *k*-mer occurrences, where *k*-mers containing an *N* are excluded.

**Figure 2.**
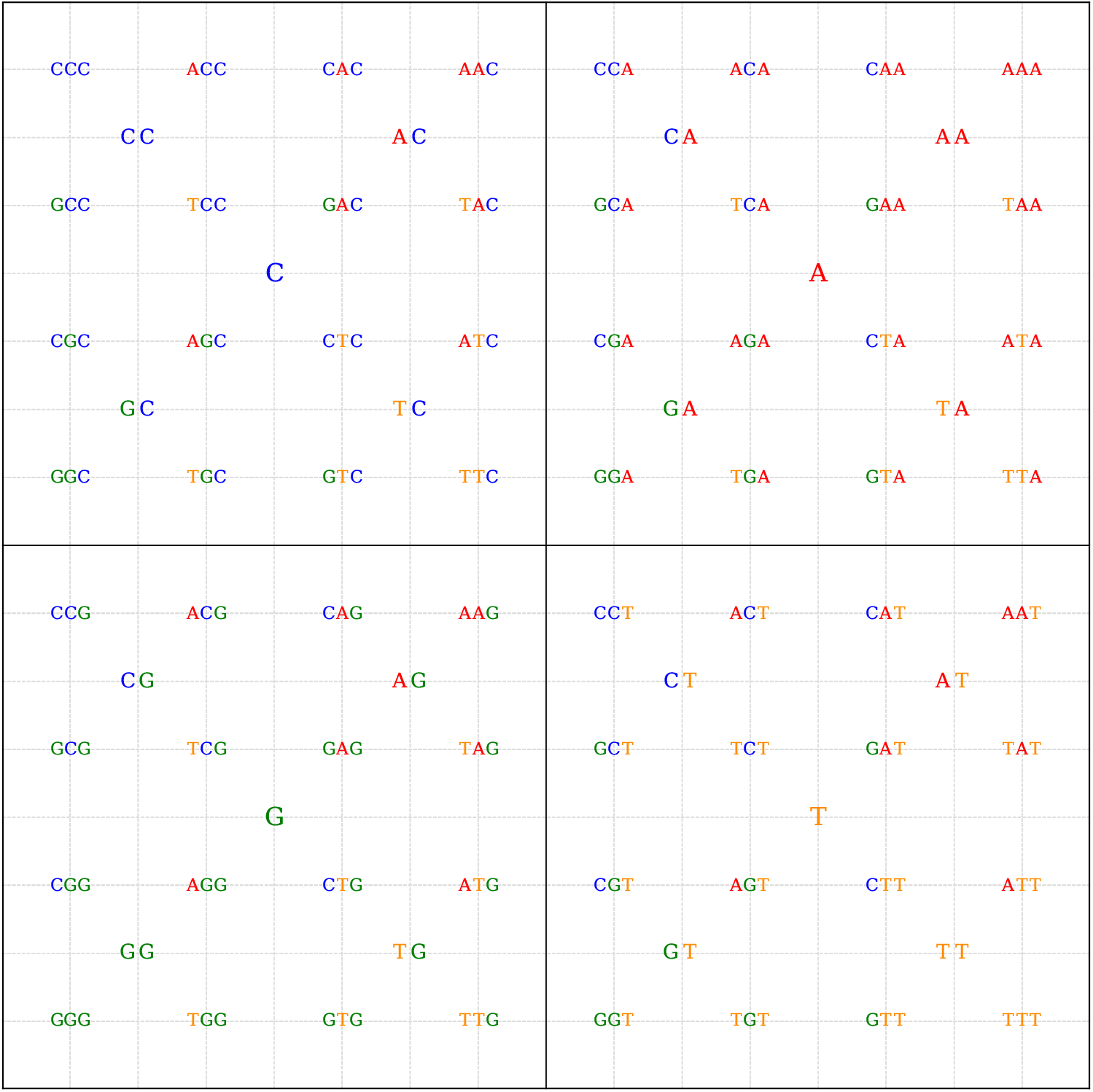
**Position of each** *k***-mer** (for *k* ∈ {1, 2, 3}) in the square [−1, 1]^2^ ⊂ ℝ^2^ given by the CGR encoding. Each nucleotide is assigned to a vertex of the square to guide the encoding of *k*-mers as in Definition 1. We can observe that a subdivision of one level of a sub-square (*i*.*e*. from *k* to *k* + 1) corresponds to adding the letters *A, C, G, T* as prefixes to the *k*-mer encoded in the center of such sub-square, following the orientation of the vertices.

**Definition 2** (Frequency matrix of CGR (FCGR)). *Let* 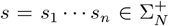 *be a sequence, and let k be an integer. Then the Frequency matrix of CGR, in short FCGR, of the sequence s is a* 2^*k*^ × 2^*k*^ *2-D matrix F* = (*a*_*i,j*_), 1 ≤ *i, j* ≤ 2^*k*^, *i, j* ∈ ℕ. *For each k-mer b* ∈ {*A, C, G, T*}^*k*^, *we have an element a*_*i,j*_ *in the matrix F, that is equal to the number of occurrences of b as a substring of s. Moreover, the position* (*i, j*) *of such element is computed as follows:*

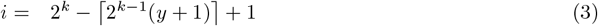

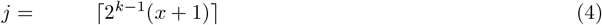

*where* (*x, y*) *is the CGR encoding for the k-mer b*.

It is immediate to obtain a 2-D image from an FCGR matrix; it suffices to rescale all its values to the interval [0, 255] (8 bits). Examples of these images for three different species are shown in Figure 1, and the ordering that *k*-mers adopt in the representation is shown in Figure 2, for *k* ∈ {1, 2, 3}, we can see that *k*-mers sharing their suffix are close in the matrix.

### 2.3 Convolutional Neural network for the FCGR

Convolutional layers [22], the building block of Convolutional Neural Networks (CNN), are commonly used to process spatially structured data, such as images. Formally, given an input tensor 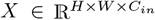, where *H* and *W* denote the height and width of the input, respectively, and *C*_*in*_ the number of input channels, a convolutional layer applies a set of *K* learnable filters 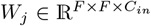, with *j* ∈ [1, *K*] and *F* known as the kernel size, to generate an output feature map 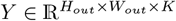. The dimensions *H*_*out*_ and *W*_*out*_ depend on the kernel size *F*, and another parameter called stride, which controls the step size of the filter as it moves across the input, in addition, can be affected if padding of the output is used across the height or width dimensions.

When the FCGR is the input tensor *X*, the filters *W*_*k*_ correspond to a 2-dimensional matrix, and we have a single channel. As we can see in Figure 1, different species show different patterns when seen through the lens of the FCGR, these patterns are the result of how *k*-mer occurrences are placed in the 2-dimensional matrix. In Figure 2 we can illustrate this ordering for *k* = 3, notice for example that in the top-left, the sub-square with center *CC* contains the 3-mers *ACC, CCC, GCC*, and *TCC*, all sharing the same 2-long suffix *CC*. More generally, given *k*, in the FCGR representation, *k*-mers sharing the *l*-long suffix (*l < k*) are contained in a sub-square of dimension 2^*k*−*l*^ × 2^*k*−*l*^ in the FCGR.

This observation is useful for using the *k*-mer ordering in the FCGR when defining a convolutional layer to exploit this ordering. We define a convolutional layer, named *ConvFCGR*, with kernel size 2^*k*−*l*^ and stride 2^*k*−*l*^ to be applied over the FCGR matrix, in this case, each convolutional filter processes only the number of occurrences of the *k*-mers sharing a given *l*-long suffix, for 0 *< l < k*.

Under the umbrella of supervised metric learning, we propose the `CNNFCGR` architecture (see Figure 3), a convolutional neural network defined by stacking *ConvFCGR* layers and whose output is an *n*-dimensional vector, the so-called *embedding*. Unlike traditional supervised learning approaches designed for mere classification into a fixed number of classes, metric learning focuses on learning distance metrics or similarity measures tailored to some specific tasks. The goal of metric learning is to guarantee that similar instances are closer in the embedding space, and dissimilar ones are further apart, clustering the input data in the n-dimensional space. This is achieved using ad-hoc loss functions, most notably contrastive loss [13], or triplet loss [38] by directly optimizing the embeddings.

**Figure 3.**
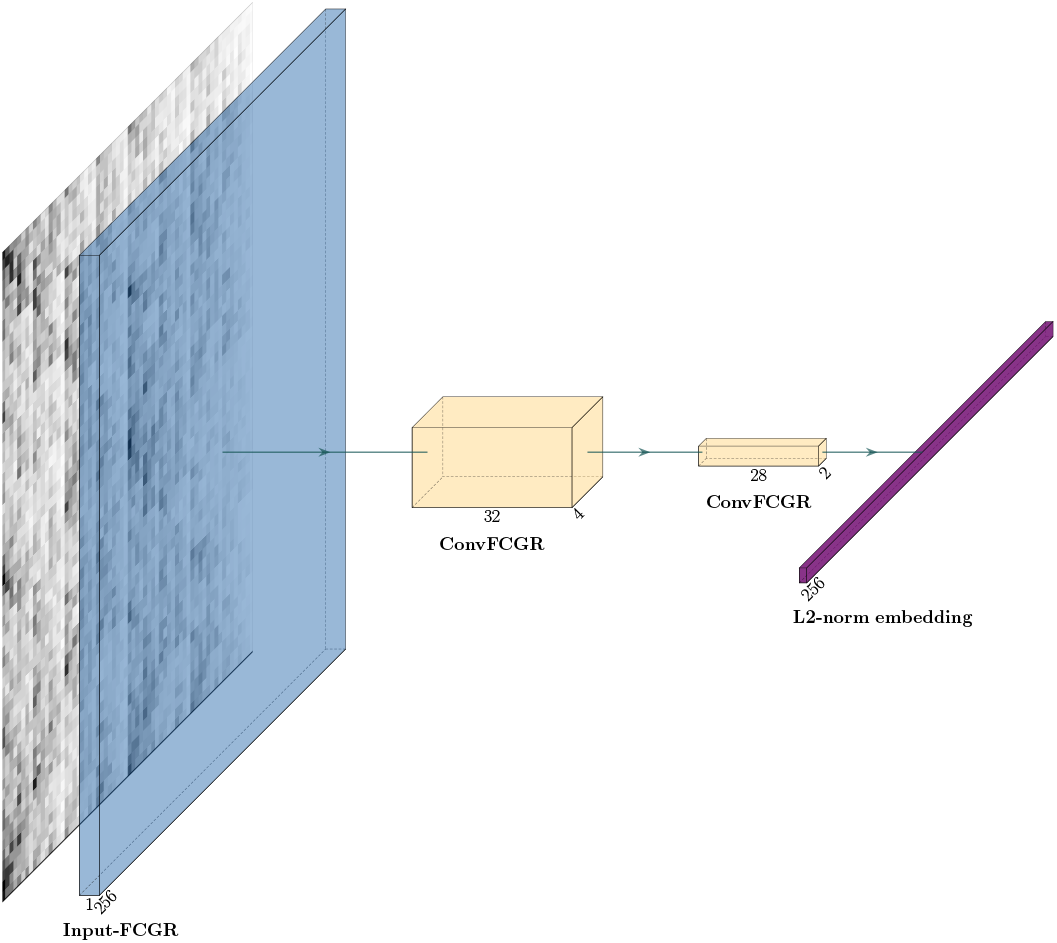
`CNNFCGR` architecture with input size 2^8^ × 2^8^ × 1, representing the FCGR of a (draft) assembly form 8-mers. The architecture is a concatenation of two *ConvFCGR* layers, one of level 2, and one of level 1, with number of filters 32 and 24, respectively. The output of the encoder is an L2 normalized *n*-dimensional vector (**Embedding**), represented by a dense layer with *n* neurons. The ReLU activation function follows all *ConvFCGR* layers, and batch normalization in the filter axis.

#### 2.3.1 ConvFCGR layer

The *ConvFCGR* layer of level *l* corresponds to a convolutional layer with kernel size 2^*l*^, stride 2^*l*^, and no padding. Hence, for an input of size 2^*k*^ × 2^*k*^, the *ConvFCGR* of level *l* outputs a tensor of dimension 2^(*k*−*l*)^ × 2^(*k*−*l*)^. Given *l* levels and *f* filters, the convolutional FCGR layer is denoted by *ConvFCGR*(*l, f*). In Figure 4 are illustrated *ConvFCGR*(1, 1) and *ConvFCGR*(2, 1) on an input of dimension 2^3^ × 2^3^.

**Figure 4.**
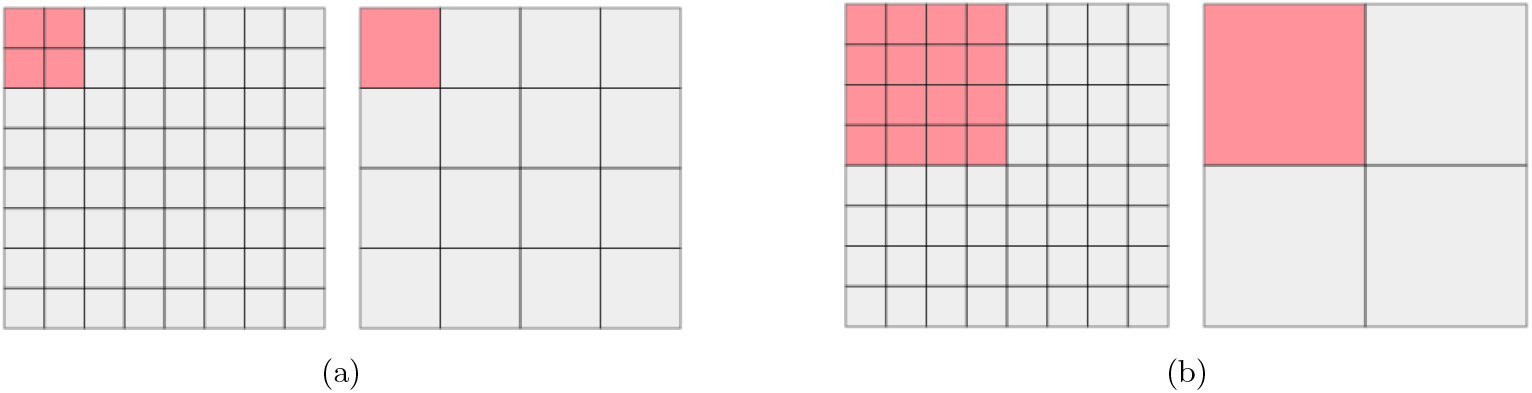
ConvFCGR layer. Gray grids correspond to the input and output of the layers. Red grids represent the position where a filter of a layer is applied. In (a) a *ConvFCGR*(1, 1) is applied to the input (FCGR) tensor of size 2^3^ × 2^3^, level *l* = 1 correspond to a layer with stride and kernel equal to 2, *i*.*e*. the output is a tensor of size 2^2^ × 2^2^. In (b) *ConvFCGR*(2, 1) is applied to the same input. This time, since *l* = 2, the layer uses 16 positions of the tensor, and outputs 1.

#### 2.3.2 Encoder: CNNFCGR architecture

Given an input tensor of dimensions (2^*k*^ × 2^*k*^ × 1) (FCGR from *k*-mers with depth channel included) the encoder is defined as a concatenation of two layers: *ConvFCGR*(2, 4 × *k*) and *ConvFCGR*(1, 4 × (*k* −1)), each followed by a batch normalization in the channel axis. This is then flattened and fed to a dense layer of *n* neurons, with an L2 normalization, so embeddings have a norm equal to 1, and Euclidean distances between embeddings lie in [0, 2]. In Figure 3 we illustrate the `CNNFCGR` architecture for 8-mers.

### 2.4 Learning embedding representation of assemblies

In what follows, we describe how the `CNNFCGR` architecture is trained. It receive as input the FCGR matrix with rescaled, and use the Rectified Adam optimizer [27] with a learning rate of 0.001, combined with the Lookahead optimizer algorithm, which improves the learning stability and lowers the variance of its inner optimizer with negligible running time and memory usage [54].

#### 2.4.1 Preprocessing of the FCGR matrix

Each FCGR was normalized to the [0, 1] interval by dividing all *k*-mer occurrence counts by the maximum value within the corresponding FCGR. In addition, we implemented an optional clipping step in which FCGR values above the *p*-th percentile are truncated prior to normalization. This procedure reduces the influence of highly overrepresented *k*-mers, thereby preventing them from dominating the normalized representation.

#### 2.4.1 Supervised Metric Learning

The goal of supervised metric learning in this study is to construct an embedding space for DNA sequences (bacterial assemblies) in which sequences from the same bacterial species are close in Euclidean distance, while sequences from different species are farther apart. The `CNNFCGR` (described in Section 2.3.2) is designed to create such embeddings from the FCGR of assemblies based on their species labels. It receives as input the FCGR matrix and outputs a *n*-dimensional vector. We use the triplet loss function [38] to learn such embedding representations.

#### 2.4.2 Embedding assemblies with the triplet loss

Given a triplet 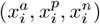, the triplet loss is defined as follows,

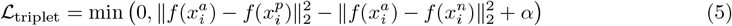

where 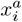 is the anchor sample, 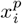 is the positive sample (same class as the anchor), 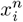 is the negative sample (different class from the anchor), *f*(·) is the embedding function that maps inputs into the feature space, ∥ · ∥_2_ is the Euclidean (L2) distance, *α* is the margin enforced between positive and negative pairs, usually defined in the interval [0.1, 0.5]. In our case, *f*(·) denotes the encoder, and the pair 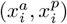 are two FCGRs from the same species, while 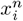 corresponds to an FCGR of a different species.

Notice that by the definition of ℒ_triplet_ not all triplets might contribute to the learning process. In fact, those triplets 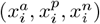 such that

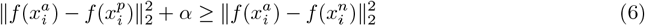

already satisfy the desired separability condition, hence ℒ_triplet_ = 0. In this case, the computation of the gradient is meaningless, therefore it does not influence the update of weights in the neural network.

As explained in [38], choosing triplets is crucial to use the triplet loss effectively. A triplet of interest (for which ℒ_triplet_ ≠ 0) can be categorized into *semi-hard* and *hard* triplets.

In this context, given an anchor 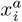, a hard-positive is computed as the argmax 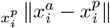, while a hard-negative is computed as the argmin 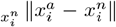. These examples are the most difficult to separate among those that violate the condition from Equation (6). Since computing the argmax and argmin is computationally expensive over the entire dataset (*e*.*g*. for a dataset of *N* elements, the number of possible triplets is *O*(*N*^3^)), in practice, these *hard* triplets are chosen from a mini-batch (the subset of data used in each iteration of an epoch). A drawback of selecting the hardest negatives is that they can lead to bad local minima early on in training [38]. On the other hand, a triplet 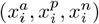 is called *semi-hard* when

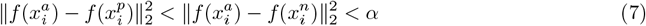

so even when 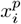 is closer to the anchor than 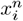, it still lies inside the margin *α*.

The suggested procedure in [38] when using the triplet loss is to start training with semi-hard triplets, and then focus on hard triplets. In our experiments we saw that the use of hard triplets (after training with semi-hard triplets) worsen the results, so we considered only semi-hard triplets for our experiments.

### 2.5 Large bacterial data sets

Our goal is to index and query large bacterial datasets. In [15] the collection of all bacterial data in the European Nucleotide Archive (ENA) available up to 2018 was consolidated into a data set of around 661K (draft) assemblies. Then it was updated with all the bacterial assemblies up to 2023, and named as AllTheBacteria [15] data set.

The assemblies in these two data sets were assembled with the same pipeline using short reads and correspond to isolated assemblies, *i*.*e*., each one of them corresponds to a single species. The main differences between these data sets are (1) how a *high-quality* assembly is defined and (2) how the assemblies are assigned a unique species label. In the 661K dataset, the abundance of species in the sample is obtained with a Kraken [51]-Bracken [28] pipeline, and the label of an isolated assembly is defined as the most abundant species in the sample if its abundance exceeds 90%. In contrast, in the AllTheBacteria data set, the assignment of a label consists of estimating the abundance of species directly from the isolated assembly using sylph [39], where the most abundant species in the assembly is assigned as its label if the abundance is greater than 99%. The number of assemblies and species for both datasets is reported in Table 2.

**Table 1.** List of tools tailored and/or adaptable for taxonomy classification in bacteria. For each tool, we point out if: it can be used for taxonomy classification, the type of query sequence, if it is a machine learning-based method (Hybrid means that part of the approach is machine learning-based), if it rely on an index (Hybrid means that part of the approach is index-based), and the minimum taxonomy level of the query results (when more than one taxonomy level is defined, it means that the tool can output either of the two depending on its confidence). `Tiara` and `EukRep` have as their main purpose the classification of eukaryotic sequences, while `MycoAI` fungal sequences, but have included or can be extended for prokaryotic taxonomy classification. `LexicMap` and `Phylign` are alignment-based tools where query results can be used for taxonomy classification. For `Taxometer`, the minimum taxonomy level depends on the dataset used to train the model. `TYGS` is provided as a web service. `HiTaxon` is an ensemble method that combines machine learning and index-based methods.

| Tool | Taxonomy Classification | Type Query | Machine Learning | Index-based method | Minimum Taxonomy Level |
| --- | --- | --- | --- | --- | --- |
| DeepMicrobes [25] | Yes | short read | Yes | No | species |
| Kraken [52] | Yes | short read | No | Yes | species |
| Kraken2 [51] | Yes | short read | No | Yes | species |
| Centrifuge [20] | Yes | short read | No | Yes | species |
| Centrifuger [42] | Yes | short and long reads | No | Yes | species |
| Taxor [44] | Yes | long read | No | Yes | species |
| HiTaxon [45] | Yes | short read | Hybrid | Hybrid | species |
| Taxometer [21] | Yes | contig | Yes | No | species |
| EukRep [48] | Yes | 5000 bp | Yes | No | domain |
| Tiara [19] | Yes | contigs | Yes | No | domain |
| MycoAI [36] | Yes | type query | Yes | No | species |
| LexicMap [40] | No | short and long reads (>500bp) | No | Yes | species |
| Phylign [4] | No | reads/contigs | No | Yes | species |
| TYGS [33] | Yes | assembly | Partial | Yes | species or subspecies |
| fIDBAC [24] | Yes | assembly | No | Yes | species |
| GAMBIT [29] | Yes | assembly | No | Yes | genus or species |
| GSearch [56] | Yes | assembly | No | Yes | species |
| PanSpace | Yes | assembly | Yes | Yes | species |

**Table 2.**
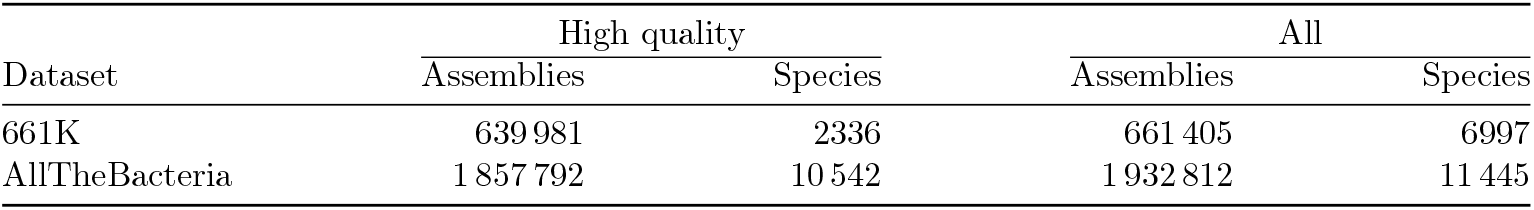
Bacterial datasets. The total number of species in the AllTheBacteria database was obtained by counting all unique values in the column label of the file sample2species2file.tsv.gz from https://ftp.ebi.ac.uk/pub/databases/AllTheBacteria/Releases/0.2/metadata/, since only the number of species of highquality assemblies was reported in the manuscript.

High-quality assemblies are defined by several criteria, based on the length of the assembly, number of contigs, N50, and the most abundant species. Under these high-quality criteria, the 96.76% and 96.12% of the assemblies are considered high-quality in the 661K and AllTheBacteria data sets, respectively.

#### 2.5.1 Mini-batch generation and training configuration

A mini-batch consists of a subset of the data set randomly sampled to be fed to a network in the training phase. For example, in the 661k bacterial data set, we find 2336 species belonging to highquality assemblies, while in the AllTheBacteria bacteria data set, the number of species reported is 10542 (see Table 2), with most of the assemblies belonging to only a small portion of the available species, *e*.*g*. the top 10 species constitute the 75% of the high-quality data set. When dealing with a highly imbalanced data set and a large number of labels, the generation of mini-batches requires special considerations, since a random selection will most likely lead to discard the underrepresented species. With the triplet loss, we must provide a way to subsample the data set for creating the triplets. For this purpose, we use a mini-batch generator that first randomly selects a portion of the labels (species), and then selects a fixed number of FCGRs for each label. Thus, each time we train the `CNNFCGR`, the triplets are created from a balanced data set with a restricted number of possible anchors.

We selected a mini-batch size of 512 assemblies, corresponding to (randomly selected) 64 species, and multiplied by a factor of 5 the number of mini-batches generated for training and validation steps, *i*.*e*. the number of mini-batches corresponds to 5 × the ratio between the FCGRs in the training set (as well in the validation set) and the mini-batch size, thus increasing the number of triplets seen during each training epoch.

We set the training of `CNNFCGR` to 100 epochs, with an early stopping of patience of 30 epochs and a patience learning rate of 15 epochs with a factor of 0.1, both monitoring the validation loss.

Training, validation, and test sets were built as follows, for each species with more than 12 samples in the data set, we randomly split them into training, validation, and test sets in proportion 80 : 10 : 10. We consider only species with this representativity since in our experiments we want to query the index (built from training and validation sets) and retrieve up to 11 neighbors, so we can be sure that there are at least 11 assemblies for each species in the index.

### 2.6 Creating the Index

We recall that our objective is to learn a function *f* : 𝒮_*k*_ → ℝ^*n*^ (for *n* ≪ 4^*k*^) that reduces the dimensionality of the *k*-mer occurrences of an assembly by encoding some relevant information in an n-dimensional vector (embedding) for its classification, where 𝒮_*k*_ represents the space of *k*-mer occurrences for the set 𝒮 of assemblies. We can approximate *f* by choosing a mapping *E* : 𝒮_*k*_ *↦* ℝ^*n*^, from now on called the *encoder*, corresponding to the trained `CNNFCGR` (see Figure 3).

The `PanSpace` index ℐ consists of three components: (1) a FAISS index ℱ [10, 18] storing embeddings for each assembly in 𝒮, (2) the encoder *E*, and (3) the labeling function *λ*. The encoder *E* generates an *n* − dimensional embedding for each assembly (*s* ∈ *𝒮*), which is stored in ℱ. FAISS provides a compact (less than 2GB for ~ 1.6 million embeddings of dimension 256), fast, and easy-to-query structure. To link embeddings to species labels via *λ*, we maintain a corresponding list of labels for all entries in ℱ. Finally, to query new assemblies against 𝒮, we apply the encoder ℐ to map them into the same embedding space, so *E* is an integral part of the index ℐ.

### 2.7 Querying the Index

A query to the index ℐ is defined as a (draft) assembly *s*_*q*_ (in fasta format, and which might not belong to S), and the query results correspond to a list with the *K*-closests (a user defined parameter) embeddings in the FAISS index ℱ and their labels given by the labeling function *λ*. We assign the label *L* to *s*_*q*_, where *L* corresponds to the most common label in the query result.

The process of going from an (possibly draft) assembly to the embedding space where the indexed assemblies lie involves (1) counting *k*-mers of the assembly, (2) creating the FCGR matrix, and (3) using the encoder *E* to get an embedding in R^*n*^, as illustrated in Figure 5.

**Figure 5.**
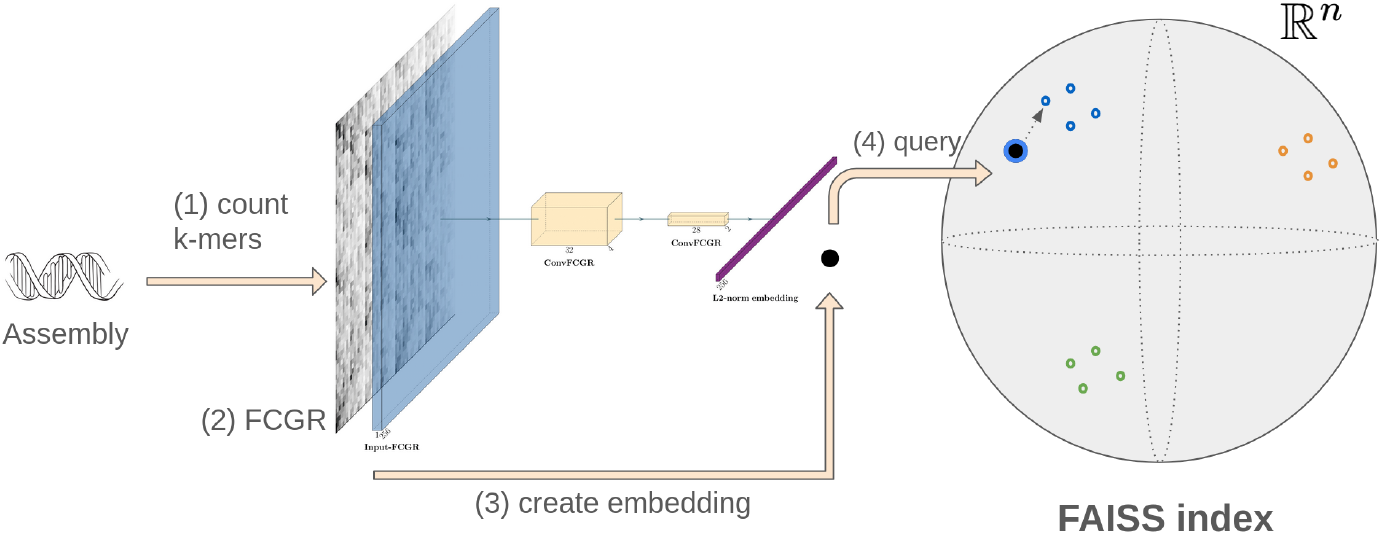
Query index. Given a query (draft) assembly, we start by (1) counting *k*-mers, (2) creating its FCGR representation, and (3) producing the embedding representation by feeding the FCGR to the Encoder. Finally, (4) we query the FAISS index to retrieve its *K*-closest neighbors and assign a label to the query assembly based on the majority of species in the query result.

## 3 Experiments

We recall that existing query systems that can employ databases of the size of the AllTheBacteria data set (see Table 2) require excessive amounts of computational resources. This hampers their widespread application due to their dependency on expensive compute clusters and huge consumption of electricity notwithstanding their excellent accuracy in classification. The current driving open challenge in prokaryote classification is to make substantial savings in terms of computational resources during all stages of classification, while preserving state-of-the-art accuracy in classification. PanSpace addresses exactly this challenge.

For this, one needs to re-consider construction, storage, and the querying of the index itself. The popular, greater vision is to run classification on laptops, as a convenient, resource-friendly and independent option. The great practical values are obvious. Quick, laptop based classification of prokaryotes will enable field workers to classify the bacterial specimens encountered immediately, without incurring delays due to (up to daylong) data transfer to compute clusters and the corresponding (excessive) expenses they can be subject to.

In view of this situation, we pursue two goals, one primary and one secondary. Our primary goal is to demonstrate that our classification system indeed introduces drastic savings in terms of time, space, and costs. The secondary goal we pursued was to demonstrate that we maintained the superior levels of accuracy in classification already reached by earlier work. Regarding our secondary goal, we note that the clearly leading and most accurate classifier is `GSearch` [56]. In the following, we therefore focus on a comparison with `GSearch` in particular. The corresponding analysis involves inspecting the ANI between assemblies and query results delivered either via the `PanSpace` or the `GSearch` index.

In the following, we exclusively consider high-quality assemblies to avoid hard-to-control biases induced by low-quality assemblies that could blur our evaluation. In addition, for technical reasons, to ensure that our evaluation is not subject to irregularities due to data set size imbalances (which may introduce fatal biases during validation, leading to suboptimal choices of hyperparameters), we consider only species sporting at least 13 different assemblies. Note that once training procedures have been approved through validation, these data set imbalances no longer matter—the only consideration remaining is to have training data sets sufficiently large.

Considering subsets of species of at least 13 assemblies ensures that at least 1 assembly can be part of the test set (at a data split training : validation : test of 80:10:10), with at least 11 assemblies for each species in the index (see additional experiments in the Supplementary Material). The size of the data set is 1 841 109 assemblies, of which 90% (1 656 597) is used to build the indexes, and 10% (184 512) for queries. For `PanSpace`, train and validation sets comprises the 90% used to build the index, and the test set correspond to the 10% used for queries.

For `PanSpace`, we conduct experiments using the `CNNFCGR` architecture (see Section 2.3) with 8-mers, applying a preprocessing step that clips values at the 80th percentile. We compare these results against `GSearch` configured with three sketching algorithms: `GSearch`-setsketch, `GSearch`probminhash, and `GSearch`-optdenminhash, all operating on 16-mers. In `GSearch`-setsketch, sketches of size 4,096 are built on the presence or absence of k-mers, whereas `GSearch`-probminhash and `GSearch`-optdenminhash use sketches of size 12,000 and incorporate k-mer frequencies.

### 3.1 Indexing the bacterial data set

Creating the `PanSpace` indexes for the AllTheBacteria data set requires to store the FCGR representations of assemblies. This amounts to 131 GB and 480 GB for 7and 8-mers, respectively. Instead, creating and storing `GSearch` indexes involves storage of the collection of the assemblies (in fa.gz, that is in compressed FASTA format), requiring 4.1 TB, more than 8.5 times than for 8-mer and more than 31 times than for 7-mer based `PanSpace` indexes.

We recall that the index that `PanSpace` generates and maintains is composed of (1) the FAISS index, which stores the embeddings of the assemblies to be indexed, (2) the *Encoder* part of the trained model, which transforms assemblies (sets of contigs, so sets of sequences) into their embeddings (real-valued vectors), and (3) the list of labels for each assembly in the index.

#### 3.1.1 Computational resources: Building the index

In Table 3, we report the sizes of the different (`PanSpace` and `GSearch`) indexes, the maximum RAM usage required during construction, as well as the time that elapses to build the indexes. The storage footprint of the Encoder itself is negligible, requiring 1.1MB of storage, while the list of labels requires only 60MB, also only a minor contribution to the space requirements. Note that storing the `PanSpace` index primarily depends on the dimension *n* of the embeddings that one uses (see Supplementary Material).

**Table 3.**
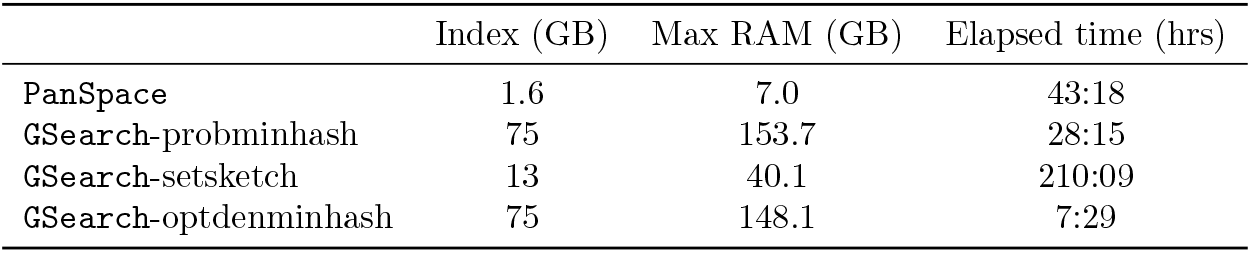
Computational resources to build the index.. The column Index corresponds to the size of the index size (disk space) comprising 1 656 597 assemblies (90% of the data set). Maximum RAM usage and Elapsed time required for building the indexes are reported in the lasts two columns.

It is also important to consider the impact of the sketch size on the resource requirements of `GSearch`. While all variants rely on 16-mers, `GSearch`-probminhash and `GSearch`-optdenminhash use sketches of size 12 000, resulting in indexes that occupy about 75 GB of disk space. In contrast, `GSearch`-setsketch operates with sketches of the much smaller size of 4 096, reducing disk usage to just 13 GB, a dramatic difference in storage demands.

Among the three modes of `GSearch`, `GSearch`-setsketch is the one requiring the least RAM (40.1 GB), while consuming the most runtime (210 hours). `GSearch`-optdenmihash and `GSearch`-probminhash, on the other hand, require ~ 150 GB of RAM, where `GSearch`-optdenminhas is 4× faster to construct than `GSearch`-probminhash, with the latter requiring 7.5 hours for construction.

In the context of accuracy in classification (see the next section for full details), `GSearch`-optdenminhash obtains optimal results. Compared to `GSearch`-optdenminhash, the `PanSpace` index is ~ 47× smaller than its counterpart and requires ~ 21× less RAM for construction. On the other hand, it requires 6 times more runtime (~ 43 vs.~ 7 hours).

`GSearch`-setsketch is second in classification. In this case, the `PanSpace` index is 8× smaller than the index of `GSearch`-setsketch and requires ~ 6× less RAM. Here, `PanSpace` is also faster, requiring 5 times less runtime than `GSearch`-setsketch (~ 43 vs.~ 210 hours).

#### 3.1.2 Computational resources: Querying

To assess the computational resources required to query the indexes built with `PanSpace` and `GSearch`, we randomly selected 1000 draft assemblies from the AllTheBacteria data set. These assemblies are from the test set; therefore, they have not contributed to constructing the index.

`GSearch` and `PanSpace` were run using 48 threads involving CPUs alone. In all cases, queries were performed on assemblies provided as compressed FASTA (fa.gz) files. In Table 4 we report both the maximum RAM usage and the time required to query the 1000 assemblies for both `PanSpace` and `GSearch`.

**Table 4.** Computational resources to query the index. We randomly selected 1000 draft assemblies from the AllTheBacteria data set (from the 10% of the data set defined as the test set) and query the indexes built with `PanSpace` and `GSearch` (built with the 90% of the data set). In all cases we used 48 threads and only CPU.

|  | Max RAM (GB) | Elapsed time (mm:ss) |
| --- | --- | --- |
| **PanSpace** | 3.7 | 2:28 |
| **GSearch-probminhash** | 84.6 | 6:03 |
| **GSearch-setsketch** | 17.1 | 15:39 |
| **GSearch-optdenminhash** | 81.2 | 5:00 |

`PanSpace` spends most of the time on FCGR generation (which includes *k*-mer counting; with `PanSpace` operating at *k* = 8). The creation of embeddings for the 1000 assemblies takes roughly 1 second in total with the `CNNFCGR` architecture.

When evaluating the most accurate version of `GSearch` against `PanSpace`, we find that `GSearch`optdenminhash requires 5 minutes with a peak memory usage of 81.2 GB, whereas `PanSpace` completes the task in only 2 minutes and 28 seconds with a maximum of 3.7 GB of RAM. This shows that `PanSpace` is approximately 10× faster and 22× more memory-efficient when querying 1,000 assemblies.

### 3.2 Accuracy of querying the index

Here, we compare the accuracy achieved by `PanSpace` and `GSearch`, when querying the respective indexes, for which we evaluated the space and time requirements in the subsection before. We recall that the test data set of assemblies not used during the construction of the indexes consists of 10% of the data set (amounting to 184 512 assemblies), while, accordingly, the indexes contain the remaining 90% of assemblies (amounting to 1 656 597 assemblies).

Given a query assembly, `PanSpace` identifies the embedding of the assembly in the index that has minimal distance to the query embedding, since in nearest-neighbor the closest neighbor represents the statistically most significant match [8]. The label assigned to a query is inherited from its closest neighbor in the index (see Section 2.7). To generate the embedding for a query assembly, the corresponding assembly is run through the encoder.

For `GSearch`, we follow the same approach as in `PanSpace`: the label for a query is assigned based on its closest neighbor in the `GSearch` index. However, in this case, the index is based on genome sketches and ANI similarity, rather than embeddings.

#### 3.2.1 Evaluation Metrics

We evaluate the performance of classification using the index-retrieval methodology with three standard metrics: Precision, Recall, and F1-score, computed for each species.

For a given species in the query set, Recall measures the proportion of query assemblies from that species that are correctly classified, relative to the total number of assemblies of that species in the query set. Precision, on the other hand, measures the proportion of correctly classified query assemblies for that species among all assemblies predicted to belong to it. The F1-score is the harmonic mean of Precision and Recall, providing a single measure that balances both metrics and is particularly useful when they differ significantly.

These metrics are especially informative in unbalanced datasets. For example, if the index always predicts the most common species, it would achieve high recall for that species (since most of its true assemblies are recovered) but low precision, as many assemblies from other species would be incorrectly labeled as such. The resulting F1-score would thus reflect this imbalance, remaining low despite the high recall.

The same metrics are also computed at the genus level, where genera represent a higher taxonomic rank grouping related species.

Because we are dealing with a multi-class classification problem on a highly imbalanced dataset (see Section 2.5), we report the macro-averaged Precision, Recall, and F1-score. The macro average corresponds to the unweighted mean of the per-class metrics, assigning equal weight to each class irrespective of the number of samples it contains. This choice avoids bias toward highly represented species, which would dominate the results under micro-averaging. Results are summarized in Table 5.

**Table 5.** Accuracy and macro average Precision, Recall and F1-score at species and genus levels, considering the predicted species as the species of the closest neighbor in the index. The best results are highlighted in **bold**.

| Classification metrics using 1-nearest neighbor |  |  |  |  |  |  |
| --- | --- | --- | --- | --- | --- | --- |
|  | Species |  |  | Genus |  |  |
|  | Precision | Recall | F1-score | Precision | Recall | F1-score |
| PanSpace | 0.993 | 0.990 | 0.991 | **0.999** | **0.999** | **0.999** |
| GSearch-probminhash | 0.992 | 0.992 | 0.992 | 0.991 | 0.992 | 0.992 |
| GSearch-setsketch | 0.995 | 0.993 | 0.994 | 0.995 | 0.993 | 0.994 |
| GSearch-optdenminhash | **0.996** | **0.995** | **0.995** | **0.999** | 0.997 | 0.998 |

At the species level, all three modes of `GSearch`, and `PanSpace` have precision, recall and f1-score above 0.99, with `GSearch`-optdenminhash being the best overall. Importantly, differences between `GSearch` and `PanSpace` only show at the third digit behind the decimal point; it is fair to rate performance in accuracy to be pretty much on a par with one another. In other words, `PanSpace` metrics come very close to the best results of `GSearch`, with a relative difference of at most 0.5% in all metrics, a negligible increase in errors. At the genus level, `PanSpace` outperforms all `GSearch` versions in terms of all metrics. This points out that `PanSpace` captures the taxonomic hierarchies better than `GSearch`, which one can consider a favorable effect, because it points out that the `PanSpace` index integrates species in a more holistic, comprehensive manner.

We also observe that the metrics for `GSearch`-probminhash at the genus level are lower than at the species level. At first glance, this may appear counterintuitive, since one might expect classification to improve when grouping species into broader taxonomic categories. However, aggregating species into genera alters both the number of classes under evaluation and their relative representation in the dataset. This shift in class distribution can reverse apparent trends in performance, an instance of the well-known Simpson’s paradox [41].

In Table 6, we present a breakdown of classification errors for each tool at both the species and genus level. Notably, we observe that `GSearch` occasionally fails to return a result for certain queries — a behavior that never occurs with `PanSpace`. In contrast, `PanSpace` consistently outputs the closest neighbors for each query, ensuring that a classification is always produced.

**Table 6.** Classification errors using 1-nearest neighbor on a test set of 184 512 assemblies. The column ‘No answer’ counts the number of query assemblies for which the index did not output a result. The column ‘Wrong Genus’ counts the mislabeled assemblies for which the genus is also incorrect. The column ‘Wrong Species/Correct Genus’ counts the mislabeled assemblies for which the genus is correct.

|  | No answer | Wrong Genus | Wrong Species/<br>Correct Genus |
| --- | --- | --- | --- |
| PanSpace | 0 | 11 | 47 |
| GSearch-probminhash | 0 | 18 | 27 |
| GSearch-setsketch | 3 | 14 | 25 |
| GSearch-optdenminhash | 0 | 14 | 27 |

Additionally, in Figure 6, we report precision, recall, and F1-score for each species, grouped into bins according to their representativeness in the index. The results clearly show that data representativeness has a major impact on performance across all tools: species with ≤100 assemblies in the index often exhibit substantially lower scores, while highly represented species (*>*10 000 assemblies) consistently achieve near-perfect classification. Notably, `GSearch`-probminhash is the only approach that struggles in the lowest representativeness bin ( ≤100), with three species showing precision, recall, and F1-scores equal to 0. These findings suggest that as more genomic data become available, index-based methods will benefit significantly from increased representativeness, leading to more accurate and robust classification.

**Figure 6.**
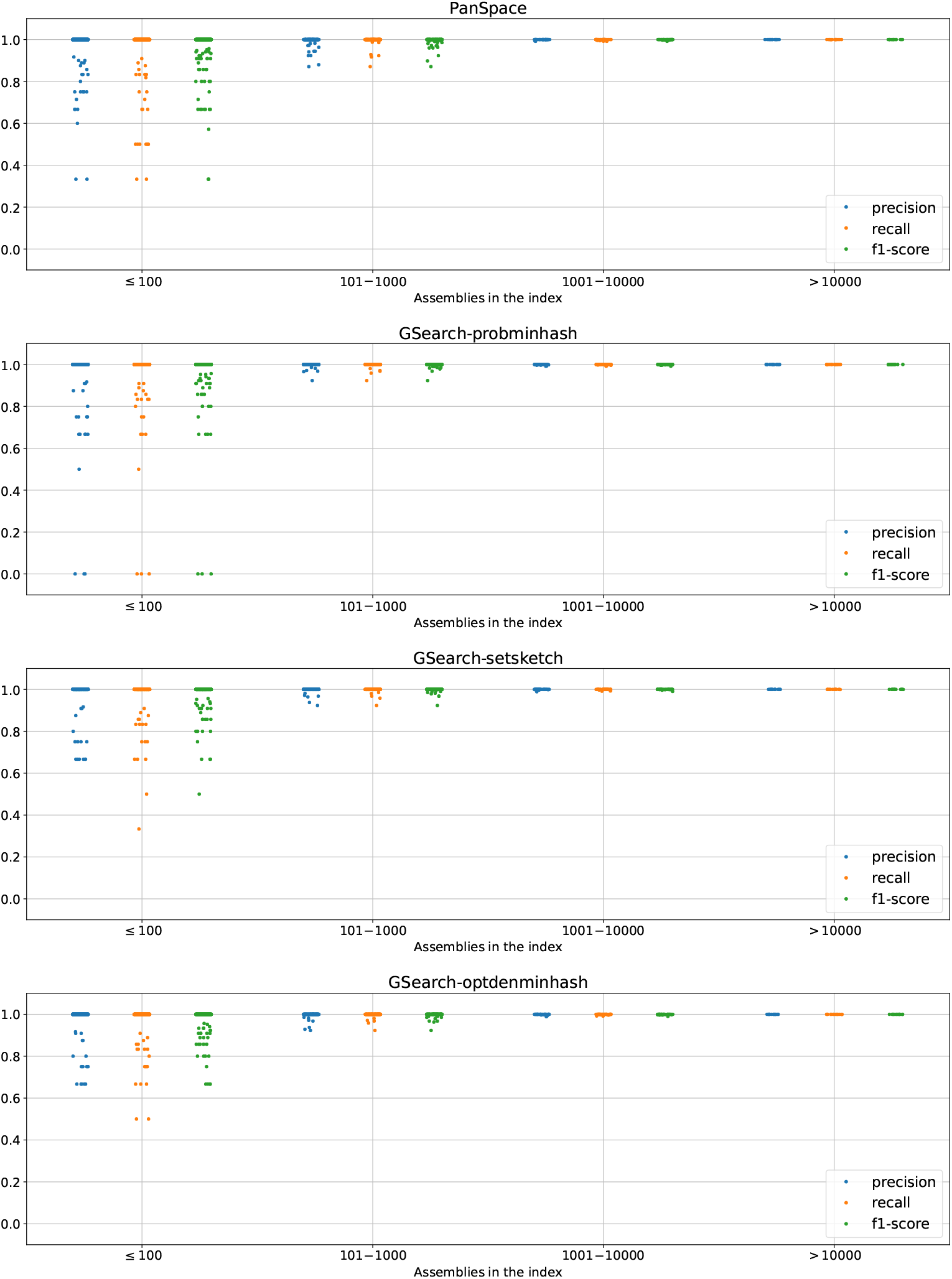
Metrics by species support in the index. We grouped the individual metrics (precision, recall, and f1-score) for each species into bins ([≤ 100, 101 −1000, 1001 −1000, *>* 10000]) showing the representativity of each species in the index. Lower representativity ( ≤ 100) impacts negatively across all tools. While higher representativity leads to perfect classification by all tools. Leading to the conclusion that while more data becomes available, index-based tools will improve.

#### 3.2.2 Embedding proximity reflects high ANI values between assemblies

In this experiment, we show that assemblies that are close w.r.t. the Euclidean distance between their embeddings obtained with `PanSpace` have a large ANI value.

We randomly selected 1000 assemblies (from the test set) from each of the 30 most represented species in the AllTheBacteria data set, agreeing with the set of assemblies selected for assessing the computational resource requirements when querying the index in Section 3.1.2. First, we query the `PanSpace` index with these 1000 assemblies, and compute the ANI value between the query assembly and each of their 11 closest neighbors (the query result) using `fastANI` v1.34 with default parameters. We perform the same query with `GSearch`. The corresponding resulting ANI values for both `PanSpace` and `GSearch` are shown in Figure 7. We observe that `GSearch` concentrates most of its ANI values close to 1, which is expected since this index explicitly relies on ANI values between assemblies for its construction. On the other hand, `PanSpace` also shows large ANI values for all queries. Importantly, the query results (in both cases) are above the ANI threshold 95, which determines that two assemblies belong to the same species. There was one case (sample-ID query SAMEA388208), for both `GSearch` and `PanSpace`, for which the closest neighbor results in an assembly with ANI distance below the ANI threshold 95.

**Figure 7.**
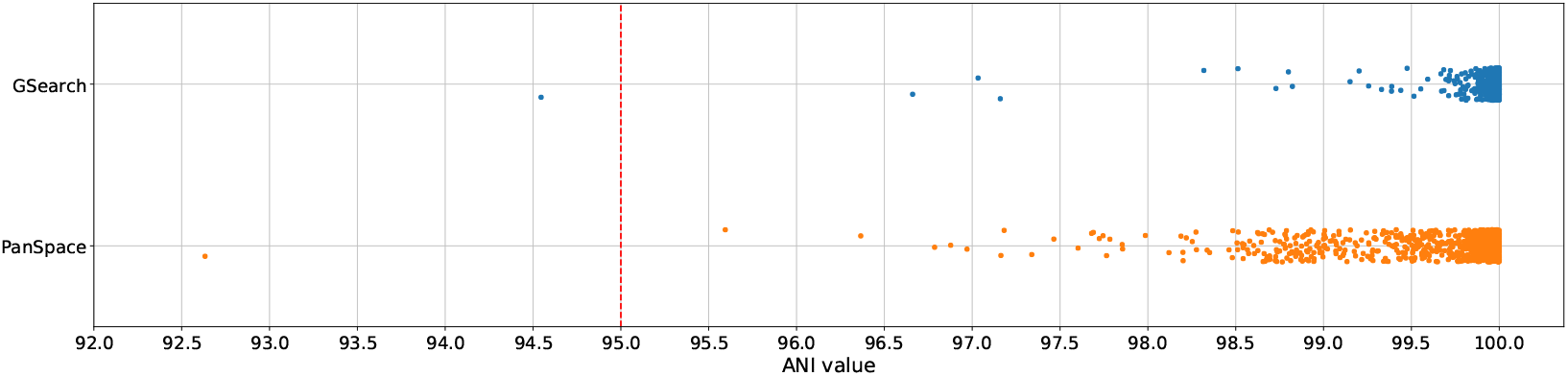
**Stripplot of ANI values** between query sequences and their closest neighbor in indexes built with `PanSpace` (orange) and `GSearch` (blue). We randomly selected 1000 draft assemblies from the AllTheBacteria data set (from the 10% of the data set defined as the test set) and query the indexes built with the 90% of the data set. We compute the ANI similarity (x-axis) between each query assembly and its closest assembly in the indexes. A vertical red dashed line indicates the 95 ANI value threshold for determining that two assemblies belong to the same species. ANI value was computed using `fastANI` v1.34 with default parameters. `GSearch`-setsketch was used in this case.

Secondly, we explore how the assemblies cluster when using their embeddings provided by `PanSpace` as the basis of the clustering. We randomly selected 25 species (2325 assemblies from the test set), and created a UMAP reduction of them [32], shown in Figure 8^3^. We observe distinct, well-separated clusters corresponding to different species. Most notably, groups of species from the same genus appear to be closer together to each other than groups of species that do not belong to the same genus.

**Figure 8.**
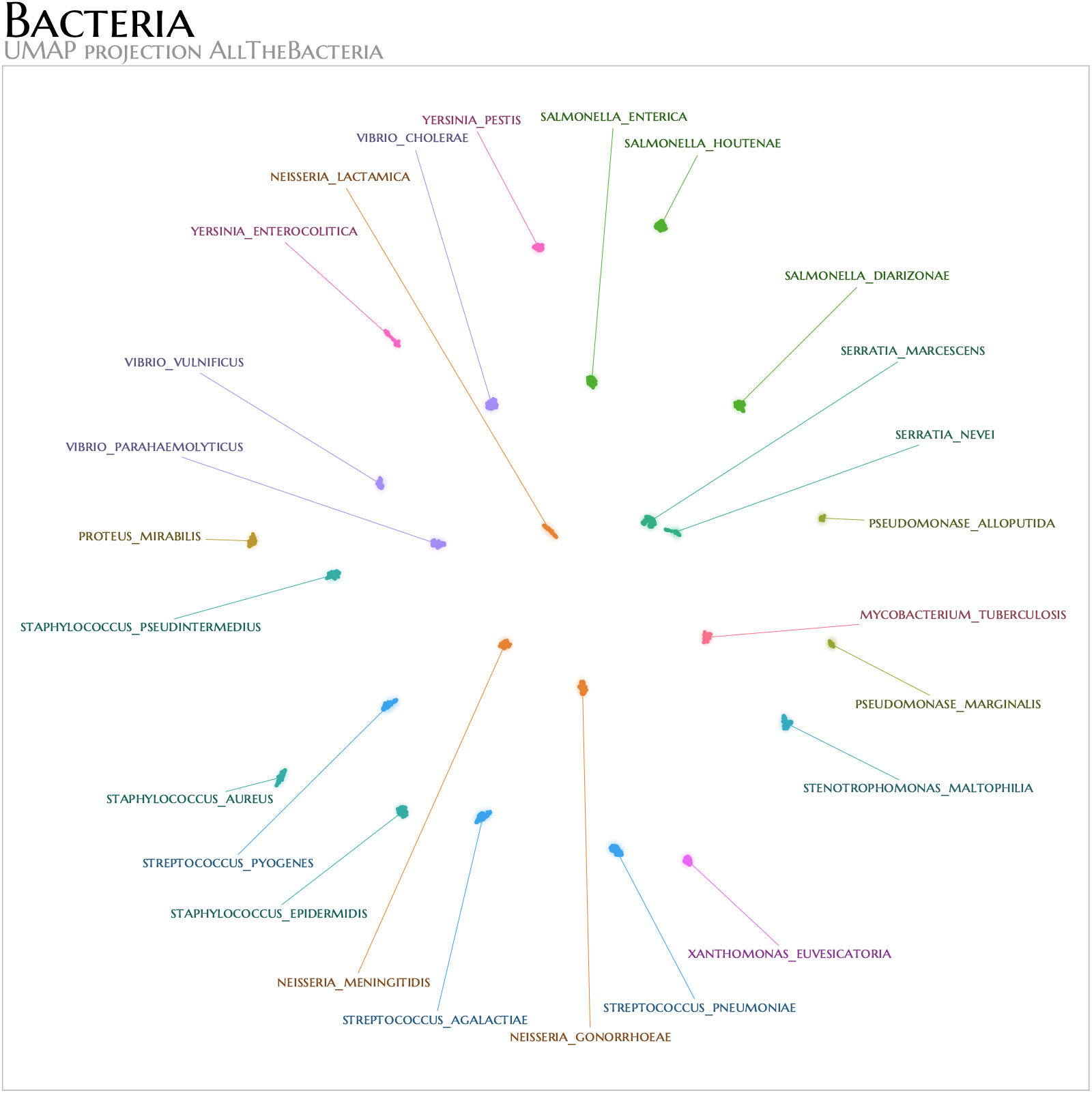
**UMAP** PanSpace. We randomly selected 25 species (2325 assemblies) in the AllTheBacteria data set (all assemblies were selected from the test set). Using the embedding representation of each assembly computed by `PanSpace` we create a UMAP projection in 2-dimensions using default parameters. Species from the same genus have identical colors.

### 3.3 Experimental Setup

All experiments for `PanSpace` were conducted on a desktop computer with the following characteristics: AMD Ryzen 7 16 cores, with 1TB of disk space, 16 GB of RAM, NVIDIA GeForce RTX 4080, using CUDA Version: 12.1. The server runs Linux as the operating system.

Experiments regarding `GSearch` (and `PanSpace` in the query performance) were conducted on a server equipped with 64 CPU cores (AMD EPYC 7301, 16-core, 2.2 GHz), 256 GB of RAM, and 9.1 TB of storage. The server runs Linux as the operating system.

The Python environment setup for training, creating, and querying the index is defined by the following libraries and versions: tensorflow v2.15.0, faiss-gpu v1.7.2, scikit-learn v1.3.2, complexcgr v0.8.0, and Python v3.10.13.

### 3.4 Data Availability

The AllTheBacteria dataset was downloaded from https://ftp.ebi.ac.uk/pub/databases/AllTheBacteria/Releases/0.2/.

Experiment results reported in this manuscript can be found in Zenodo at the following link https://zenodo.org/records/14936601.

FCGR dataset were uploaded to HugginFace at the link https://huggingface.co/datasets/koke/AllTheBacteria-FCGR-8mer

`PanSpace` is avaialable under the GNU 3.0 license at https://github.com/pg-space/panspace.

## 4 Discussion

We have introduced `PanSpace`, a deep learning tool designed to (1) index large-scale databases of bacterial genome assemblies and (2) query these indexes to identify the species from which genomic sequences of unknown origin stem. In particular, we have indexed the currently largest database available, AllTheBacteria, resulting in an index requiring less than 2 GB of disk space. `PanSpace` has also been very efficient in constructing the index, at a peak memory usage of only 7 GB and requiring only 43 hours for construction. This is very favorable in comparison with the leading state-of-the-art tool `GSearch`, which requires 13 GB to store the index in its most space-efficient version (`GSearch`-setsketch), which takes more than 40 GB of RAM and nearly 9 days for construction. Faster versions that `GSearch` can offer, taking only up to seven and a half hours for construction, so being 5-6 times faster than `PanSpace`, all yield indexes of 75 GB, that is, nearly 47 times larger than that of `PanSpace`, and require nearly 150 GB of RAM for their construction, that is, more than 21 times more RAM than `PanSpace` requires. While the trade-off between space and time is already very favorable for `PanSpace` anyway, it is important to realize that space requirements are the currently most pressing constraints when operating by means of devices of smaller size, such as laptops.

It is important to keep in mind that databases like AllTheBacteria are steadily growing due to the rapid accumulation of genome assemblies. This aggravates the situation if classification tools do not operate at a possible minimum of resource requirements. In summary, `PanSpace` has proven to be substantially more resource friendly than the current state of the art, reducing resource requirements by an order of magnitude or more. Given the rapid growth of databases, this holds great promise for the near- and mid-term future.

Of course, we have also evaluated the accuracy of `PanSpace` in terms of the classification of query sequences, compared to the leading state-of-the-art tool `GSearch`, which exhibited outstanding performance rates when predicting the origin of the query sequences at the level of species and at the level of genuses. We have demonstrated that `PanSpace` preserves the splendid performance rates at the level of species, and even exceeds the performance rates at the level of genuses. Beyond just ensuring that `PanSpace` also exhibits performance rates of utmost excellence, its superiority at the level of genuses indicates that the `PanSpace` embeddings capture the evolutionary hierarchy inherent to the corresponding taxonomies better than earlier approaches. UMAP based visualizations corroborate this intuition.

In addition, our results have shown that the assemblies in the database with which the query assemblies were matched agree with the approved standards that underlie the identification of species. The assemblies in the database with which `PanSpace` has matched the query assemblies have consistently exceeded an average nucleotide identity (ANI) of 95, which reflects a widely accepted threshold for two sequences that are to belong to identical species. Visualizations of the `PanSpace` embeddings (via UMAP) reveal that they form clearly delimited clusters at the level of species, where the clusters of species are positioned close to each other if they belong to the same genus. The latter analysis confirms that `PanSpace` has captured the taxonomic hierarchies inherent to prokaryotic assemblies better than the earlier approaches.

In summary, `PanSpace` has introduced an efficient deep-learning framework for indexing and querying large-scale bacterial genome collections. It achieves comparable or superior accuracy to `GSearch` while requiring less time, memory, and storage. Beyond query performance, the embeddings learned by `PanSpace` capture meaningful biological structure, forming species-level clusters and reflecting genus-level relationships. These results demonstrate not only the robustness of `PanSpace` as a practical tool for large-scale genome analysis but also its potential as a foundation for future methods leveraging learned embeddings for comparative genomics and microbial taxonomy.

From the point of view of methodical advances, `PanSpace` has integrated data structures that facilitate the use of the already abundant and plentiful knowledge about evolutionary taxonomies in the context of a deep learning based framework. In this, the decisive factors of `PanSpace` have been to, first, harness the power of deep learning in terms of dealing with abundant knowledge and second, to organize the data that allows to extract the knowledge in a way that accounts both for the biological meaning inherent to the data and their rapid processing through year-long, fine-tuned and therefore ultra-fast techniques that specialize in the analysis of image type data structures.

`PanSpace` has been based on the so-called Frequency Chaos Game Representation (FCGR) that transforms genomic squences into two-dimensional patterns that preserve biological contexts in an approved manner, namely in terms of *k*-mers. It is important to know that small subsequences usually reflect sequential patterns that play crucial roles in regulatory mechanisms. The FCGR arranges *k*-mers into an image type structure, which, subsequently, can be exploited in convolutional layer based neural networks. This has been opening up the wide range of (often ultra-fast) techniques that have been available for convolutional neural networks in the more recent past.

Another key point of `PanSpace`‘s success has been the integration of a specialized data loader that addresses the strong class imbalance inherent to large-scale bacterial datasets. This data loader mitigates the issue via balanced sampling at the level of species. As a result, one can ensure the appropriate integration of also unterrepresented species into systematic training procedures. This in turn improves on generalization across both common and rare species, a key factor in achieving consistent performance.

Equally importantly, `PanSpace`‘s index consists solely of an array of multidimensional vectors and the Encoder that transforms (FCGR based) species into such vectors. The underlying metric learning architecture has been extremely lightweight compared to other possible architectures that can serve this purpose. In summary, `PanSpace` has proven to be substantially more resource-friendly than earlier approaches, and is even suitable to run the corresponding large-scale classification tasks even on personal computers and embedded systems.

We have taken a decided step towards harnessing the power of deep learning when classifying genomic sequences in terms of their evolutionary origin. In that, we have outlined a systematic, extremely accurate and, in particular, ultra resource-friendly protocol. Importantly, this protocol lays the flexible foundation for extensions of all kinds, when it comes to the classification of taxa.

Of course, there are various opportunities to extend and also improve our work. An obvious issue from which one can elaborate further is the imbalance of the origin the sequences provided for training originate. For example, usually, newly discovered bacterial species suffer from being underrepresented in the training data. Recent advances in metric learning, such as the Threshold Consistency Margin (TCM) loss [55] have the potential to mitigate the issue by providing a basis for indexing strategies that are robust, adaptable, and improve generalization.

Another avenue that is worth being considered is the construction of indexes that facilitate the classification of shorter sequences, such as reflecting raw read data or single contigs. The integration of the classification of shorter sequences will open up the wide field of metagenomics in terms of the evolutionary classification of the environments the metagenomes are drawn from at incomplete or without any prior assembly of the sequenced reads. One can achieve this at no sacrifice in terms of assembly-based classification by employing AI techniques that support the processing of multiple data types or modalities. Various such techniques have recently been presented in the literature, spearheaded by examples that support the integration of text and images into common spaces.

Last but not least, recent work on the evaluation of the importance of features supports the evaluation of the embedded sequences in terms of the identification of subsequences that drive classification more than others. In other words, the integration of techniques that highlight the regions in the sequences that reflect evolutionary hotspots will lead to an improved understanding of evolution itself. Because understanding evolution is the key driver of biological research, the integration of such techniques connects us back to the fundamentally most driving questions.

## 6 Acknowledgments

This research work has received funding from the European Union’s Horizon 2020 Research and Innovation Staff Exchange programme under the Marie Sklodowska-Curie grant agreement No. 872539 (Pangaia) and from the research and innovation programme under the Marie Sklodowska-Curie grant agreement No 956229 (Alpaca). This research work is also supported by the grant MIUR 2022YRB97K, Pangenome Informatics: from Theory to Applications (PINC). We also thank the team behind the AllTheBacteria data set that made possible the execution of this work.

## Footnotes

1 gisaid.org ACCESSION DATE 03-03-2025

2 https://tygs.dsmz.de/

3 Image created with https://datamapplot.readthedocs.io

