## Supplementary Material for "PanSpace: Fast and Scalable Indexing for Massive Bacterial Databases"

[1]DISCo, Univ. Milano – Bicocca, Viale Sarca, Milano, 20126, Italy
[2]Department of Applied Informatics, Faculty of Mathematics, Physics and Informatics, Comenius University in Bratislava, Mlynská dolina F1, Bratislava, 84248, Slovakia
[3]Genome Data Science, Faculty of Technology, Bielefeld University, Bielefeld, Germany
[*]first author
[†]joint last authors

November 5, 2025

#### 1 Experiments

We recall that highly accurate query systems referencing databases of the size of the AllTheBacteria data set already exist—any additional advantages in terms of classification accuracy can only be minor. However, the corresponding query systems require excessive amounts of computational resources. Therefore, these systems do not cater to adequate democratization (because of the reliance on expensive compute clusters) and sustainability (because their excessive demand of electricity).

The remaining, driving open challenge in prokaryote classification is to make substantial savings in terms of computational resources during all stages of classification, without incurring notable losses in terms of classification accuracy. The corresponding savings require to re-consider construction, storage, and the querying of the index itself. The popular and greater vision is to run classification on laptops. This will enable field workers to classify the specimens encountered swiftly, without any delay induced by (up to daylong) data transfer to compute clusters, and without being subject to excessive expenses.

In view of this situation, we pursue two goals, one primary and one secondary. Our primary goal is to demonstrate that our classification system indeed introduces massive savings in terms of time, space, and costs. Our secondary goal is to demonstrate that we maintain the superior levels of accuracy in classification already reached by earlier work. As for our secondary goal, we note that the clearly leading, most accurate classifier is `GSearch`. In the following, we therefore focus on a comparison with `GSearch` in particular. The corresponding analysis involves inspecting the ANI between assemblies and query results delivered either via the `PANSPACE` or the `GSearch` index.

In the following, we exclusively consider high-quality assemblies to avoid hard-to-control biases induced by low-quality assemblies that could blur our evaluation. In addition, for technical reasons, to ensure that our evaluation is not subject to irregularities due to data set size imbalances (which may introduce fatal biases during validation, leading to suboptimal choices of hyperparameters), we consider only species sporting at least 13 different assemblies. Note that once training procedures have been approved through validation, these data set imbalances no longer matter—the only consideration remaining is to have training data sets sufficiently large.

Considering subsets of species of at least 13 assemblies ensures that at least 1 assembly can be part of the test set (at a data split training : validation : test of 80:10:10), with at least 11 assemblies for each species in the index. The size of the data set is 1841109 assemblies, of which 90% (1656597) is used to build the indexes, and 10% (184512) for queries. For `PANSPACE`, train and validation sets comprises the 90% used to build the index, and the test set correspond to the 10% used for queries.

We provide experiments using the `CNNFCGR` architecture with 7-mers and 8-mers, and include experiments using the `ResNet50` [2] architecture, a well-known model for image classification in deep learning, as a baseline for our CNNFCGR architecture. GSearch v0.3.1 was run with three different sketching algorithms and the following parameters:

1. GSearch-probminhash:

```
gsearch --pio 1000 --nbthreads 48 tohnsw -d <dir_sequences> \
            -s 12000 -k 16 --ef 1600 -n 128 --algo prob
```

2. GSearch-setsketch:

```
gsearch --pio 1000 --nbthreads 48 tohnsw -d <dir_sequences> \
    -s 4096 -k 16 --ef 1600 -n 128 --algo hll --scale_modify_f 0.25
```

3. GSearch-optdenminhash:

```
gsearch --pio 2000 --nbthreads 48 tohnsw -d <dir_sequences> \
    -s 12000 -k 16 --ef 1600 -n 128 --algo optdens --scale_modify_f 0.25
```

#### 1.1 Indexing the bacterial data set

Creating the PANSPACE indexes for the AllTheBacteria data set requires to store the FCGR representations of assemblies. This amounts to 131 GB and 480 GB for 7- and 8-mers, respectively. Instead, creating and storing GSearch indexes involves storage of the collection of the assemblies (in fa.gz, that is in compressed FASTA format), requiring 4.1 TB, more than 8.5 times than for 8-mer and more than 31 times than for 7-mer based PANSPACE indexes.

We recall that the index that PANSPACE generates and maintains is composed of (1) the FAISS index [1, 3], which stores the embeddings of the assemblies to be indexed, (2) the *Encoder* part of the trained model, which transforms assemblies (sets of contigs, so sets of sequences) into their embeddings (real-valued vectors), and (3) the list of labels for each assembly in the index.

##### 1.1.1 Computational resources: Building the index

In Table 1, we report the sizes of the different (PANSPACE and GSearch) indexes, the maximum RAM usage required during construction, as well as the time that elapses to build the indexes. Additionally, we report the size of the Encoder for PANSPACE; we mention that the list of labels requires solely 60MB, which makes only a minor contribution to the space requirement accounts.

While the storage footprint of the encoder itself is negligible (see second column in Table 1); 0.8 GB for $n = 128$, 1.6 GB for $n = 256$, and 3.2 GB for $n = 512$), storing the PANSPACE index primarily depends on the dimension $n$ of the embeddings that is used.

It is important to note that GSearch uses 16-mers for creating sketches, because the index crucially depends on sketch size. Both GSearch-probminhash and GSearch-optdenminhash use a sketch size of 12 000, which translates into both indexes asking for 75GB of disk space. Instead, GSearch-setsketch uses a sketch size of 4096, which requires 13GB of disk space, dramatically less than when operating at sketch sizes of 12 000.

Building the PANSPACE index scales with $k$, the length of $k$-mers (because $k$ determines the dimension of the FCGRs) and $n$, as the dimension of the space of the embeddings, because the dimension $n$ has an influence on amounts of weights as part of the *Encoder* and the FAISS index. Minimum RAM usage and fastest construction time were achieved on 7-mers and embeddings of size $n = 128$, at 4.4 GB of RAM and 8 hours of runtime required. When using 8-mers and embeddings of size $n = 512$, runtime increases to 62 hours, at 11.8 GB of RAM required. Among the three versions of GSearch, setsketch is the one requiring the least RAM (40.1 GB), while consuming the most runtime (210 hours). GSearch-optdenmihash and GSearch-probminhash, on the other hand, both require ∼150 GB of RAM, while being 4× faster, that is, requiring 7.5 hours for construction (approximately matching runtime requirements for PANSPACE).

Putting the different options of both approaches into context with optimal accuracy in classification (see next section for full details), we note that optimal results for PANSPACE are obtained using 8-mers (so $k = 8$) and embedding size $n = 256$ while running GSearch in setsketch mode, on the other hand. In this case, the PANSPACE index is $8\times$ smaller than the GSearch index, and requires $\sim 6\times$ less RAM, and 5 times less runtime (42 vs. 210 hours).

Note that the ResNet50 encoder is $84\times$ larger than the CNNFCGR encoder, considerably affecting training times. However, as we will see in the next section, using the ResNet50 encoder does not lead to better results, which renders considerations with respect to ResNet50 by and large obsolete—FCGR's paired with ordinary CNN's appear to clearly lead in all disciplines.

| | Index (GB) | Encoder (MB) | Max RAM (GB) | Elapsed time (hrs) |
|---|---|---|---|---|
| CNNFCGR-$128^7$ | 0.8 | 0.6 | 4.4 | 8:04 |
| CNNFCGR-$256^7$ | 1.6 | 1.1 | 6.9 | 14:29 |
| CNNFCGR-$512^7$ | 3.2 | 2.1 | 11.0 | 16:16 |
| CNNFCGR-$128^8$ | 0.8 | 0.6 | 4.5 | 28:55 |
| CNNFCGR-$256^8$ | 1.6 | 1.1 | 7.0 | 43:18 |
| CNNFCGR-$512^8$ | 3.2 | 2.1 | 11.8 | 62:33 |
| ResNet50-$256^7$ | 1.6 | 93 | 7 | 114:40 |
| GSearch-probminhash | 75 | - | 153.7 | 28:15 |
| GSearch-setsketch | 13 | - | 40.1 | 210:09 |
| GSearch-optdenminhash | 75 | - | 148.1 | 7:29 |

Table 1: **Computational resources to build the index.**. The column Index corresponds to the size of the index size (disk space) constructed from 90% of the data set (encompassing training and validation splits), comprising 1656597 assemblies. The column Encoder corresponds to disk space claimed by each architecture trained. Embedding size is indicated next to each architecture (CNNFCGR or ResNet50), while $k$-mer size is displayed as superscript. The size of the PANSPACE index size does neither depend on $k$-mer size, nor on choice of architecture, but exclusively depends on the size of the embedding space. This explains why CNNFCGR-$256^7$, CNNFCGR-$256^8$ and ResNet50-$256^7$ share the same index size. Max RAM and Elapsed time required for building the indexes are reported in the lasts two columns. The embedding size is next to each model. The superscript represent the $k$-mer size.

###### 1.1.2 Computational resources: Querying

To assess the computational resources required to query the indexes built with PANSPACE and GSearch, we randomly selected 1000 draft assemblies from the AllTheBacteria data set. These assemblies are from the test set; therefore, they do not make part of the index.

GSearch and PANSPACE were run using 48 threads involving CPUs only. In all cases, queries were performed on assemblies provided as compressed FASTA (fa.gz) files.

In Table 2 we report both maximum RAM usage and time required to query the 1000 assemblies for both PANSPACE and GSearch. For PANSPACE, we report results using the CNNFCGR architecture with embedding size 128, 256 and 512, using 7-mers and 8-mers. For completion, we also include ResNet50 architecture with embedding size 256 using 7-mers.

In PANSPACE, most of the time is spent in FCGR generation (which includes $k$-mer counting). The creation of embeddings for the 1000 assemblies takes roughly 1 second in total with the CNNFCGR architecture (all embedding sizes) and 22 seconds with the ResNet50 architecture. In all cases, maximum RAM does not exceed 5.4GB, and requires no more than 3 minutes to complete.

In GSearch, the fastest version was GSearch-optdenminhash, requiring 5 minutes, at a maximum RAM usage of more than 80GB. Similarly, GSearch-probminhash, requires 6 minutes and 84.6GB of RAM. GSearch-setsketch is slowest, taking more than 15 minutes to complete, while requiring less RAM than the other versions, namely 17.1GB.

##### 1.2 Accuracy of querying the index

Here, we compare the accuracy achieved by PANSPACE and GSearch, when querying the respective indexes, for which we evaluated space and time requirements in the subsection before. We recall that

| | Max RAM (GB) | Elapsed time (mm:ss) |
|---|---|---|
| CNNFCGR-128$^7$ | 2.9 | 1:52 |
| CNNFCGR-256$^7$ | 3.7 | 2:12 |
| CNNFCGR-512$^7$ | 5.4 | 2:39 |
| CNNFCGR-128$^8$ | 2.9 | 2:10 |
| CNNFCGR-256$^8$ | 3.7 | 2:28 |
| CNNFCGR-512$^8$ | 5.4 | 3:00 |
| ResNet50-256$^7$ | 3.8 | 2:53 |
| GSearch-probminhash | 84.6 | 6:03 |
| GSearch-setsketch | 17.1 | 15:39 |
| GSearch-optdenminhash | 81.2 | 5:00 |

Table 2: **Computational resources to query the index**. We randomly selected 1000 draft assemblies from the AllTheBacteria data set (from the 10% of the data set defined as the test set) and query the indexes built with PANSPACE and GSearch (built with the 90% of the data set). For PANSPACE the reported results were computed using 7-mers and 8-mers. In all cases we used 48 threads and only CPU. The embedding size is next to each model. The superscript represent the $k$-mer size.

the test data set of assemblies not used during construction of the indexes consists of 10% of the data set (amounting to 184 512 assemblies), while, accordingly, the indexes contain the remaining 90% of assemblies (amounting to 1 656 597 assemblies).

Given a query assembly, for PANSPACE, we retrieve the $K$ assembly embeddings that are closest to the query, as measured per Euclidean distance varying $K$ by choosing one of $K \in \{1, 5, 11\}$. The label that we assign to the query is the one found most common among the $K$ assemblies retrieved (see **??**). For generating the embedding for the query assembly, we run the corresponding assembly through the encoder

We report results for our CNNFCGR encoders at embedding sizes $n = 128, n = 256$ and $n = 512$, both for 7-mers and 8-mers (so $k = 7$ and $k = 8$), and for the ResNet50 encoders at embedding size $n = 256$ using 7-mers ($k = 7$); these choices of parameters $k, n$ match optimal choices for CNNFCGR)

For , we assign the label to the query as per GSearch's default mode of operation, for each of the 3 different sketching algorithms, as introduced earlier.

Let us recall that PANSPACE is trained using species as labels, while GSearch relies on computing the ANI similarity from sketches. It is also worth noting that we do not provide explicit genus information during the training in PANSPACE, and in the case of GSearch, addressing genus level classification can be approached at the amino acid level (after predicting genes), using the AAI (Average Aminoacid Identity) [4]. However, we skip this step since this is not the main goal of our experiment, rather, we would like to understand if misclassified assemblies at the level of species fall into the correct genus with these approaches.

The optimal results for both GSearch and PANSPACE are achieved when using a single neighbor, whereas increasing the number of neighbors leads to a decrease in performance metrics.

### References


[1] Matthijs Douze, Alexandr Guzhva, Chengqi Deng, Jeff Johnson, Gergely Szilvasy, Pierre-Emmanuel Mazaré, Maria Lomeli, Lucas Hosseini, and Hervé Jégou. The faiss library. 2024.

[2] Kaiming He, Xiangyu Zhang, Shaoqing Ren, and Jian Sun. Deep residual learning for image recognition. In *Proceedings of the IEEE conference on computer vision and pattern recognition*, pages 770–778, 2016.

[3] Jeff Johnson, Matthijs Douze, and Hervé Jégou. Billion-scale similarity search with GPUs. *IEEE Transactions on Big Data*, 7(3):535–547, 2019.

[4] Jianshu Zhao, Jean Pierre Both, Luis M Rodriguez-R, and Konstantinos T Konstantinidis. Gsearch: ultra-fast and scalable genome search by combining k-mer hashing with hierarchical navigable small world graphs. *Nucleic Acids Research*, 52(16):e74–e74, 2024.

| | | **Macro average** | | | | | |
|---|---|---|---|---|---|---|---|
| | | Species | | | Genus | | |
| Neighbors | Model | Precision | Recall | F1-score | Precision | Recall | F1-score |
| 1 | CNNFCGR-128$^7$ | 0.945 | 0.927 | 0.930 | 0.998 | 0.998 | 0.998 |
| | CNNFCGR-256$^7$ | 0.982 | 0.967 | 0.971 | 0.997 | 0.998 | 0.998 |
| | CNNFCGR-256$^7$-clip90 | 0.983 | 0.973 | 0.975 | **0.999** | **0.999** | **0.999** |
| | CNNFCGR-512$^7$ | 0.975 | 0.967 | 0.967 | 0.996 | **0.999** | 0.997 |
| | CNNFCGR-128$^8$ | 0.939 | 0.910 | 0.916 | 0.996 | 0.995 | 0.995 |
| | CNNFCGR-256$^8$ | 0.978 | 0.965 | 0.967 | 0.998 | 0.995 | 0.996 |
| | CNNFCGR-256$^8$-clip80 | 0.993 | 0.990 | 0.991 | **0.999** | **0.999** | **0.999** |
| | CNNFCGR-256$^8$-clip90 | 0.993 | 0.987 | 0.988 | 0.999 | 0.999 | 0.999 |
| | CNNFCGR-256$^8$-mask | 0.955 | 0.932 | 0.937 | 0.995 | 0.988 | 0.990 |
| | CNNFCGR-256$^8$-mask-clip90 | 0.986 | 0.981 | 0.981 | 0.998 | 0.998 | 0.998 |
| | CNNFCGR-512$^8$ | 0.965 | 0.947 | 0.951 | 0.998 | **0.999** | 0.998 |
| | ResNet50-256$^7$ | 0.528 | 0.472 | 0.475 | 0.880 | 0.771 | 0.795 |
| | GSearch-probminhash | 0.99194 | 0.99231 | 0.99161 | 0.99148 | 0.99184 | 0.99163 |
| | GSearch-setsketch | **0.995** | 0.993 | **0.994** | 0.995 | 0.993 | 0.994 |
| | GSearch-optdenminhash | 0.994 | **0.994** | 0.993 | 0.995 | 0.994 | 0.994 |
| 5 | CNNFCGR-128$^7$ | 0.938 | 0.905 | 0.914 | 0.993 | 0.991 | 0.992 |
| | CNNFCGR-256$^7$ | 0.966 | 0.946 | 0.951 | 0.993 | 0.994 | 0.993 |
| | CNNFCGR-256$^7$-clip90 | 0.972 | 0.957 | 0.960 | 0.993 | 0.994 | 0.993 |
| | CNNFCGR-512$^7$ | 0.950 | 0.936 | 0.939 | 0.991 | 0.993 | 0.992 |
| | CNNFCGR-128$^8$ | 0.923 | 0.885 | 0.885 | 0.991 | 0.986 | 0.988 |
| | CNNFCGR-256$^8$ | 0.953 | 0.933 | 0.937 | 0.992 | 0.987 | 0.989 |
| | CNNFCGR-256$^8$-clip80 | 0.989 | 0.987 | 0.987 | 0.992 | **0.995** | 0.993 |
| | CNNFCGR-256$^8$-clip90 | 0.987 | 0.982 | 0.982 | **0.994** | 0.994 | **0.994** |
| | CNNFCGR-256$^8$-mask | 0.931 | 0.887 | 0.899 | 0.993 | 0.972 | 0.980 |
| | CNNFCGR-256$^8$-mask-clip90 | 0.982 | 0.973 | 0.975 | 0.993 | 0.994 | 0.993 |
| | CNNFCGR-512$^8$ | 0.957 | 0.922 | 0.932 | 0.993 | 0.990 | 0.991 |
| | ResNet50-256$^7$ | 0.547 | 0.495 | 0.496 | 0.887 | 0.793 | 0.816 |
| | GSearch-probminhash | 0.988 | 0.988 | 0.987 | 0.986 | 0.985 | 0.985 |
| | GSearch-setsketch | 0.992 | **0.992** | 0.991 | **0.994** | 0.992 | 0.992 |
| | GSearch-optdenminhash | **0.993** | **0.992** | **0.992** | **0.994** | 0.990 | 0.991 |
| 11 | CNNFCGR-128$^7$ | 0.898 | 0.866 | 0.873 | 0.993 | 0.986 | 0.989 |
| | CNNFCGR-256$^7$ | 0.943 | 0.926 | 0.928 | 0.992 | 0.992 | 0.992 |
| | CNNFCGR-256$^7$-clip90 | 0.958 | 0.942 | 0.944 | 0.993 | **0.994** | 0.993 |
| | CNNFCGR-512$^7$ | 0.934 | 0.915 | 0.918 | 0.990 | 0.990 | 0.989 |
| | CNNFCGR-128$^8$ | 0.894 | 0.858 | 0.865 | 0.987 | 0.980 | 0.982 |
| | CNNFCGR-256$^8$ | 0.922 | 0.905 | 0.907 | 0.987 | 0.984 | 0.984 |
| | CNNFCGR-256$^8$-clip80 | 0.985 | 0.982 | 0.982 | **0.994** | **0.994** | **0.994** |
| | CNNFCGR-256$^8$-clip90 | 0.979 | 0.972 | 0.972 | 0.993 | **0.994** | 0.993 |
| | CNNFCGR-256$^8$-mask | 0.868 | 0.819 | 0.831 | 0.961 | 0.936 | 0.945 |
| | CNNFCGR-256$^8$-mask-clip90 | 0.970 | 0.953 | 0.956 | 0.992 | 0.993 | 0.992 |
| | CNNFCGR-512$^8$ | 0.922 | 0.889 | 0.897 | 0.991 | 0.985 | 0.987 |
| | ResNet50-256$^7$ | 0.550 | 0.496 | 0.499 | 0.880 | 0.784 | 0.808 |
| | GSearch-probminhash | 0.982 | 0.984 | 0.982 | 0.982 | 0.982 | 0.982 |
| | GSearch-setsketch | **0.987** | **0.988** | **0.986** | 0.990 | 0.988 | 0.988 |
| | GSearch-optdenminhash | **0.987** | 0.987 | 0.985 | 0.990 | 0.986 | 0.987 |

Table 3: **Macro average Precision, Recall and F1-score** at species and genus levels. The embedding size is next to each model. The superscript represent the $k$-mer size. The first column (Neighbors)correspond to the number of neighbors used to compute the metrics. The first and second best results are highlighted in **bold** and with an underline, respectively.
